# MetaCoAG: Binning Metagenomic Contigs via Composition, Coverage and Assembly Graphs

**DOI:** 10.1101/2021.09.10.459728

**Authors:** Vijini Mallawaarachchi, Yu Lin

## Abstract

Metagenomics binning has allowed us to study and characterize various genetic material of different species and gain insights into microbial communities. While existing binning tools bin metagenomics de novo assemblies, they do not make use of the assembly graphs that produce such assemblies. Here we propose MetaCoAG, a tool that utilizes assembly graphs with the composition and coverage information to bin metagenomic contigs. MetaCoAG uses single-copy marker genes to estimate the number of initial bins, assigns contigs into bins iteratively and adjusts the number of bins dynamically throughout the binning process. Experimental results on simulated and real datasets demonstrate that MetaCoAG significantly outperforms state-of-the-art binning tools, producing more high-quality bins than the second-best tool, with an average median F1-score of 88.40%. To the best of our knowledge, MetaCoAG is the first stand-alone binning tool to make direct use of the assembly graph information. MetaCoAG is available at https://github.com/Vini2/MetaCoAG.

---

The development of high-throughput sequencing technologies has paved the way for metagenomics studies to analyze microbial communities without the need for culturing, especially in large scale metagenomics studies such as the Human Microbiome Project^1^. These microbial communities consist of a large number of micro-organisms including bacteria. Samples obtained directly from the environment can be sequenced to obtain large amounts of sequencing reads. In order to characterize the composition of a sample and the functions of the microbes present, we perform *metagenomics binning* where we cluster sequences into bins that represent different taxonomic groups^2^.

Next-generation sequencing (NGS) technologies such as Illumina allow us to sequence microbial communities and obtain highly accurate short sequences called *reads*. These reads can be binned^3–9^ prior to assembly, but results can be less reliable due to their short lengths^10^. Hence, a widely used pipeline for metagenomics analysis is to first assemble reads into longer sequences called *contigs* and then bin these assembled contigs into groups that belong to different taxonomic groups^2^. Current contig-binning approaches fall into two broad categories^11^: (1) *reference-based* binning approaches^7, 12–14^ which classify contigs into known taxonomic groups by comparing against a reference database and (2) *reference-free* binning approaches which cluster contigs into unlabeled bins based on genomic features of these contigs. Reference-free binning approaches^2^ have become more popular as they enable the identification of new species that are not available in the current databases. Reference-free contig-binning tools mainly make use of two features to perform binning: (1) *composition*, obtained as normalized frequencies of oligonucleotides of length *k* (referred to as *k*-mers) and (2) *coverage*, considered as the average number of reads that map to each base of the contig. These tools achieve improved performance in binning contigs by combining both the composition and the coverage information. However, it still remains challenging for these binning tools to accurately reconstruct microbial genomes of species with similar composition and coverage profiles.

Another challenge in metagenomics binning is to estimate the number of species present in a given sample. Recent binning tools have made use of *single-copy marker genes* to estimate the number of species. These single-copy marker genes appear only once in a bacterial genome and are conserved in the majority of bacterial genomes^15–17^. Hence, the presence of single-copy marker genes can be used to estimate the genome completeness and level of purity of bins. In tools such as MaxBin/MaxBin2^17, 18^, only one marker gene is utilized to estimate the number of initial bins which may lead to an underestimation of the number of species. Hence, it is worth investigating how to make use of multiple single-copy marker genes together to obtain a better estimate for the number of bins and to explore more features of contigs that can improve the binning result.

Contigs are obtained by assembling reads into longer sequences, and there are many tools to perform assembly. Most existing metagenomic assemblers^19–21^ use *assembly graphs* as the key data structure (*e.g.*, simplified de Bruijn graph^22^) to assemble reads into contigs. Previous studies indicated that contigs connected to each other in the assembly graph are more likely to belong to the same taxonomic group^23, 24^. Although popular metagenomic assemblers such as metaSPAdes^21^ output contigs along with their connection information in the assembly graph, most existing binning tools ignore the valuable connection information between contigs. More recently, tools such as GraphBin^24^, GraphBin2^25^, METAMVGL^26^ and STRONG^27^ have been developed to refine existing binning results and resolve strains using assembly graphs. These tools rely upon the bins produced by an existing binning tool and cannot dynamically adjust the number of bins. Although these tools achieve improved binning performance, they still require an initial binning result obtained from other existing binning tools and thus cannot be directly applied to bin contigs. Hence, there is a need for a stand-alone contig-binning tool that makes use of the connection information found in the assembly graph.

In this paper, we introduce MetaCoAG, a reference-free stand-alone approach for binning metagenomic contigs. In addition to composition and abundance information, MetaCoAG also makes use of the connectivity information from assembly graphs to bin contigs. More specifically, MetaCoAG estimates the number of initial bins from frequency histogram plots of all single-copy marker genes, assigns contigs into bins iteratively and adjusts the number of bins dynamically through graph-matching algorithms, and bins the remaining contigs using a label propagation method based on the assembly graph. To the best of our knowledge, MetaCoAG is the first stand-alone contig-binning tool to make direct use of the assembly graph information. We benchmark MetaCoAG against state-of-the-art contig-binning tools using simulated and real datasets. The experimental results show that MetaCoAG significantly outperforms other contig-binning tools, *e.g.*, improving the completeness of bins while maintaining high purity levels and producing more high-quality bins.

## Results

### Overview of MetaCoAG Workflow

Figure 1 shows the overall workflow of MetaCoAG. A preprocessing step (step 0 in Fig. 1) is carried out to assemble the reads into contigs and obtain the assembly graph. Metagenomic assemblers first use graph models to connect overlapping reads or *k*-mers and infer contigs as non-branching paths. After graph simplification, the vertices represent contigs and edges represent connections between contigs in the assembly graph.

**Figure 1.**
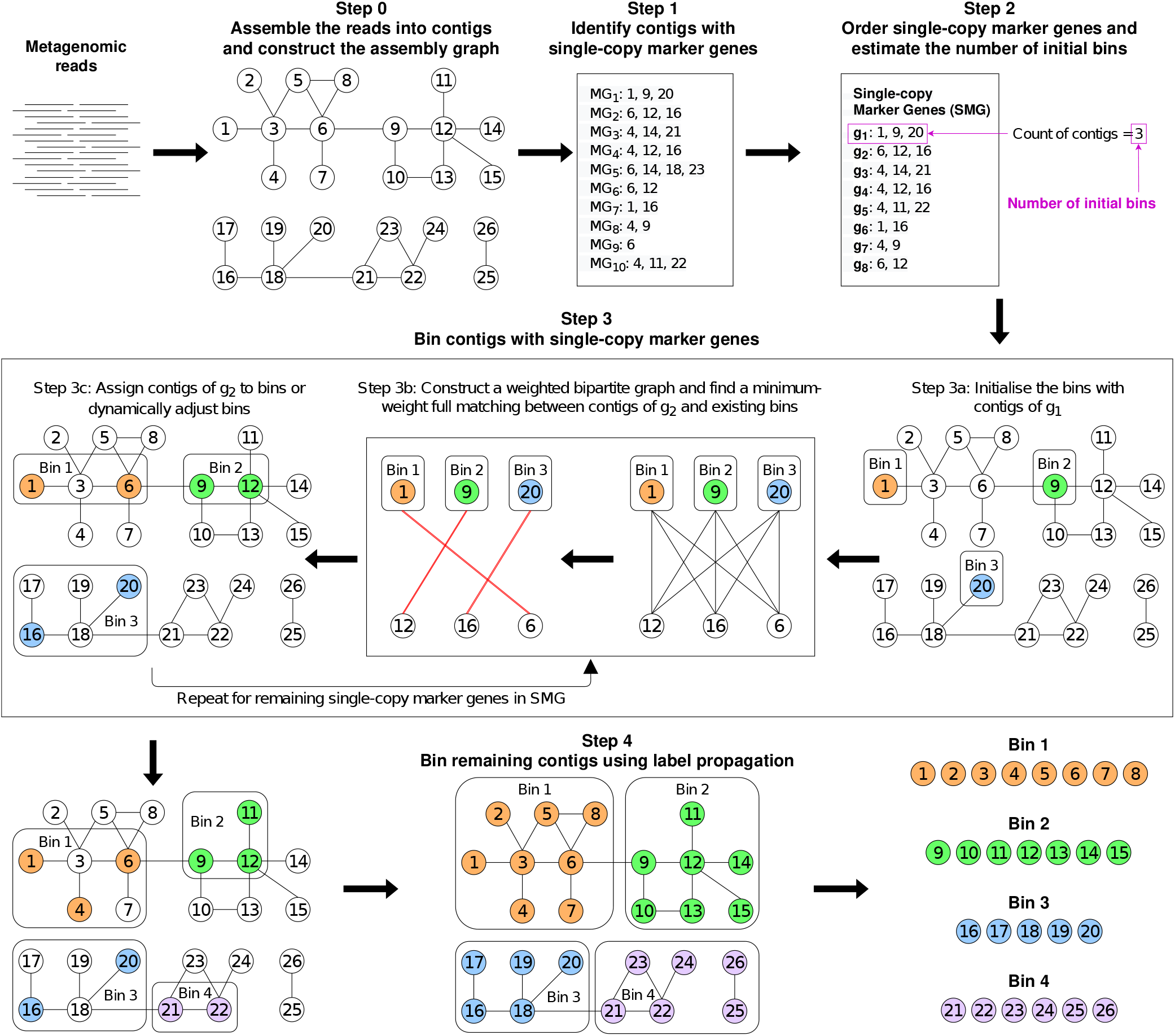
MetaCoAG Workflow. The assembly graph with contigs are provided as inputs to MetaCoAG. MetaCoAG first identifies a list of contigs that contain each single-copy marker gene. MetaCoAG further counts the number of contigs containing each single-copy marker gene and estimates the initial number of bins. Next, MetaCoAG applies a graph-matching algorithm to assign contigs that contain single-copy marker genes into bins iteratively and adjust the number of bins dynamically. Then, MetaCoAG bins the remaining contigs using label propagation algorithms based on the assembly graph and performs a postprocessing step. Finally, MetaCoAG outputs the bins along with their corresponding contigs.

Similar to previous approaches^17, 18, 28^, MetaCoAG uses 107 single-copy marker genes to distinguish contigs belonging to different species. MetaCoAG first identifies a list of contigs that contain each single-copy marker gene (step 1 in Fig. 1). MetaCoAG further counts the number of contigs containing each single-copy marker gene and estimates the initial number of bins (step 2 in Fig. 1). Then, MetaCoAG applies a graph-matching algorithm to assign contigs that contain single-copy marker genes into bins iteratively and adjust the number of bins dynamically (step 3 in Fig. 1). Finally, MetaCoAG bins the remaining contigs using label propagation algorithms based on the assembly graph (step 4 in Fig. 1), performs a postprocessing step, and outputs the bins along with their corresponding contigs. Each step of MetaCoAG is explained in detail in the Methods section.

### Benchmarks using simHC+ Dataset

We first benchmarked MetaCoAG against two popular contig-binning tools, MaxBin2^18^ and MetaBAT2^29^ on the simulated dataset **simHC+**^17^ which consists of 100 bacterial genomes (please refer to Supplementary Data 1 Table 1 for further details of the simHC+ dataset) ^1^. We evaluated the binning results of the simHC+ dataset produced by all the tools using the two popular evaluation tools AMBER^31^ and CheckM^32^. AMBER assesses the quality of bins based on the ground truth annotations provided and CheckM assesses the quality of bins based on sets of single-copy marker genes. We analyzed the purity, completeness and F1-score of the binning results calculated by AMBER (at the nucleotide level) and CheckM. MetaCoAG has recovered bins with a better trade-off between purity and completeness when compared to other binning tools (Fig. 2 (a)) with an average purity of 91.07% and an average completeness of 82.73% from AMBER and an average purity of 97.55% and an average completeness of 87.17% from CheckM. This better trade-off is demonstrated from the best F1-score results produced by MetaCoAG with a median F1-score of 95.69% from AMBER (Fig. 2 (b)) and a median F1-score of 98.48% from CheckM (Fig. 2 (c)) when compared with other binning tools. Even though MetaBAT2 has recorded the highest average purity (98.30% from AMBER and 100.0% from CheckM), it has a very low average completeness (13.02% from AMBER and 29.59% from CheckM) because all contigs shorter than 1,500bp (*i.e.* 60.49% of the contigs in the entire dataset) were discarded. Please refer to Supplementary Data 1 Table 2 for the exact values of the AMBER and CheckM results of the simHC+ dataset. We also used CheckM to count the number of high-, medium- and low-quality bins produced by all the binning tools for the simHC+ dataset (Supplementary Data 1 Table 6). MetaCoAG has recovered the highest number of high-quality bins (69 bins) and the lowest number of low-quality bins (13 bins) for the simHC+ dataset.

**Table 1.** High-quality species found from the GTDB-Tk annotations of MetaCoAG, MaxBin2 and Vamb for the real metagenomic datasets. We annotated all the high-quality bins of the real metagenomic datasets produced from MetaCoAG, MaxBin2 and Vamb using GTDB-Tk up to the species level. Then we determined whether these taxonomic groups are actually present in the original analysis. The species were determined by the classification string produced by GTDB-Tk up to species level. **✓** denotes that the species is present and **✗** denotes that the species is absent in the result/analysis. Green colored items match the original analysis whereas the red colored items do not match the original analysis. ^*^ For the COPD dataset, the species were determined as present in the original analysis based on the 50 most abundant genera presented. ^†^ These species were added to NCBI taxonomy in year 2020^37^ which is after the COPD analysis^35^.

| Dataset | Species | MaxBin2<sup>18</sup> | Vamb<sup>30</sup> | MetaCoAG | Present in original analysis |
| --- | --- | --- | --- | --- | --- |
| Sharon<sup>34</sup> | Cutibacterium avidum | ✓ | ✓ | ✓ | ✓ |
|  | Enterococcus faecalis | ✓ | ✓ | ✓ | ✓ |
|  | Peptoniphilus lacydonensis | ✓ | ✓ | ✓ | ✓ |
|  | Staphylococcus aureus | ✓ | ✓ | ✓ | ✓ |
|  | Staphylococcus epidermidis | ✓ | ✓ | ✓ | ✓ |
|  | Staphylococcus hominis | ✓ | ✗ | ✓ | ✓ |
|  | Leuconostoc citreum | ✗ | ✗ | ✓ | ✓ |
| COPD<sup>35</sup>* | Peptostreptococcus sp. | ✓ | ✓ | ✓ | ✓ |
|  | SR1 bacterium human oral taxon HOT-345 | ✓ | ✓ | ✓ | ✗<sup>†</sup> |
|  | Prevotella pallens | ✗ | ✓ | ✓ | ✓ |
|  | Haemophilus sputorum | ✗ | ✓ | ✓ | ✓ |
|  | Herbaspirillum huttiense | ✗ | ✓ | ✓ | ✓ |
|  | Capnocytophaga gingivalis | ✓ | ✗ | ✓ | ✓ |
|  | Capnocytophaga leadbetteri | ✓ | ✗ | ✓ | ✓ |
|  | Lancefieldella sp000564995 | ✓ | ✗ | ✓ | ✓ |
|  | Actinomyces graevenitzii | ✓ | ✗ | ✗ | ✓ |
|  | Actinomyces oris | ✓ | ✗ | ✗ | ✓ |
|  | Anaeroglobus micronuciformis | ✓ | ✗ | ✗ | ✗ |
|  | Eubacterium sulci | ✗ | ✗ | ✓ | ✓ |
|  | Prevotella shahii | ✗ | ✗ | ✓ | ✓ |
|  | Prevotella histicola | ✗ | ✗ | ✓ | ✓ |
|  | Lachnospiraceae bacterium oral taxon 096 | ✗ | ✗ | ✓ | ✗<sup>†</sup> |

**Figure 2.**
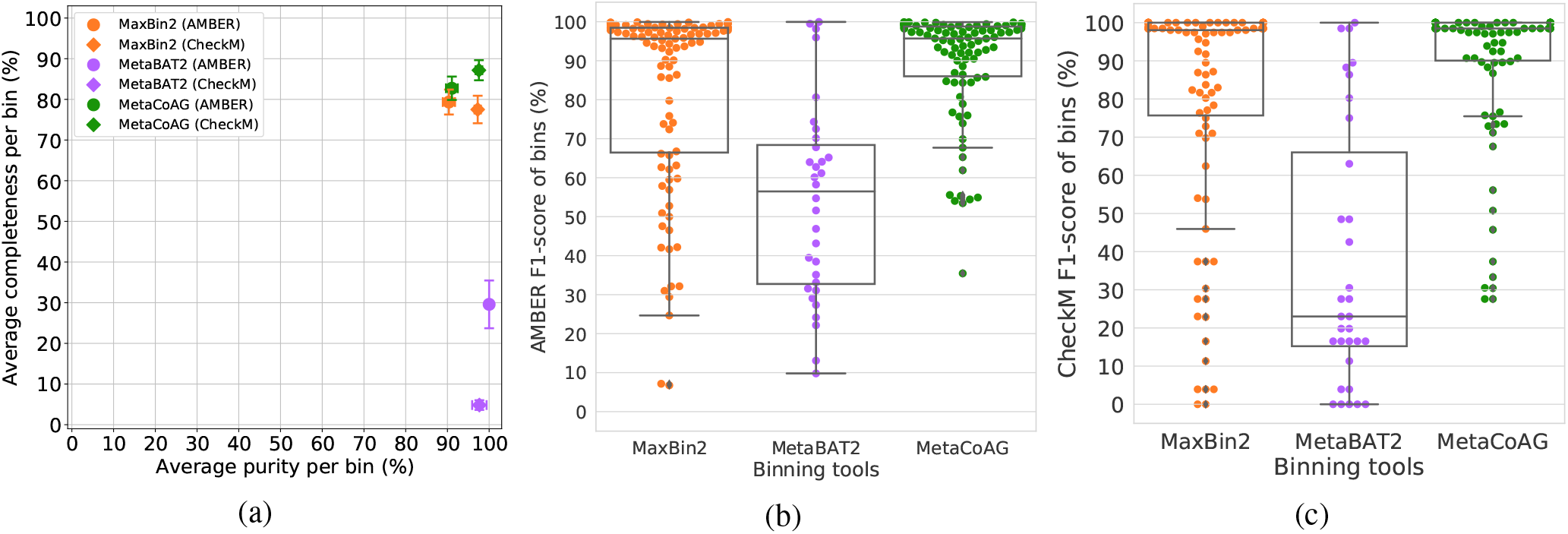
AMBER and CheckM results of the bins of the simHC+ dataset. (a) Quality of bins in terms of average completeness per bin vs. average purity per bin obtained from AMBER and CheckM. (b) F1-score of the bins obtained from AMBER. (c) F1-score of the bins obtained from CheckM.

We further used AMBER to analyze the species recovered by each binning tool for the simHC+ dataset. Out of the 100 species, MetaCoAG was able to recover more species than other tools (Supplementary Data 1 Table 4), thanks to its adaptable bin-breaking mechanism that allows to separate more species rather than combining them together. We also analyzed the F1-score of these recovered species (Supplementary Data 1 Fig. 2), and observed that MetaCoAG has recovered more species with high F1-score than the other binning tools (Please refer to Supplementary Data 1 Table 4 for comparison of the F1-score of the species recovered by MaxBin2, MetaBAT2 and MetaCoAG). Many existing binning tools assume that the oligonucleotide composition and coverage are conserved across the genome. Hence it is challenging for such tools to bin species with high variance in oligonucleotide composition and/or coverage. Moreover, these tools face difficulties when recovering species with low abundance due to the rare occurrence of species-specific signals. In Fig. 3, we visualize and compare the binning results of MaxBin2 and MetaCoAG ^2^ against the ground truth for the following species, *Pseudomonas putida* and *Arthrobacter arilaitensis*. The species *Pseudomonas putida* has a high variance in oligonucleotide composition (standard deviation > 0.015 for the tetranucleotide composition of its contigs) and thus MaxBin2 has split this species into multiple bins incorrectly (refer to Fig. 3 (a)). The species *Arthrobacter arilaitensis* has a high variance in genome coverage (standard deviation > 50× for the coverages of its contigs) and thus MaxBin2 has mis-binned some high-coverage contigs into other species with high coverage (refer to Fig. 3 (b)). However, MetaCoAG has been able to recover these species with high F1-score values, *e.g.,* improving the F1 score for *Pseudomonas putida* from 59.78% to 99.56% and improving the F1-score for *Arthrobacter arilaitensis* from 97.65% to 98.99%. Despite the high variance in oligonucleotide composition and coverage, MetaCoAG has been able to recover these species accurately, thanks to the additional connectivity information from the assembly graph.

**Figure 3.**
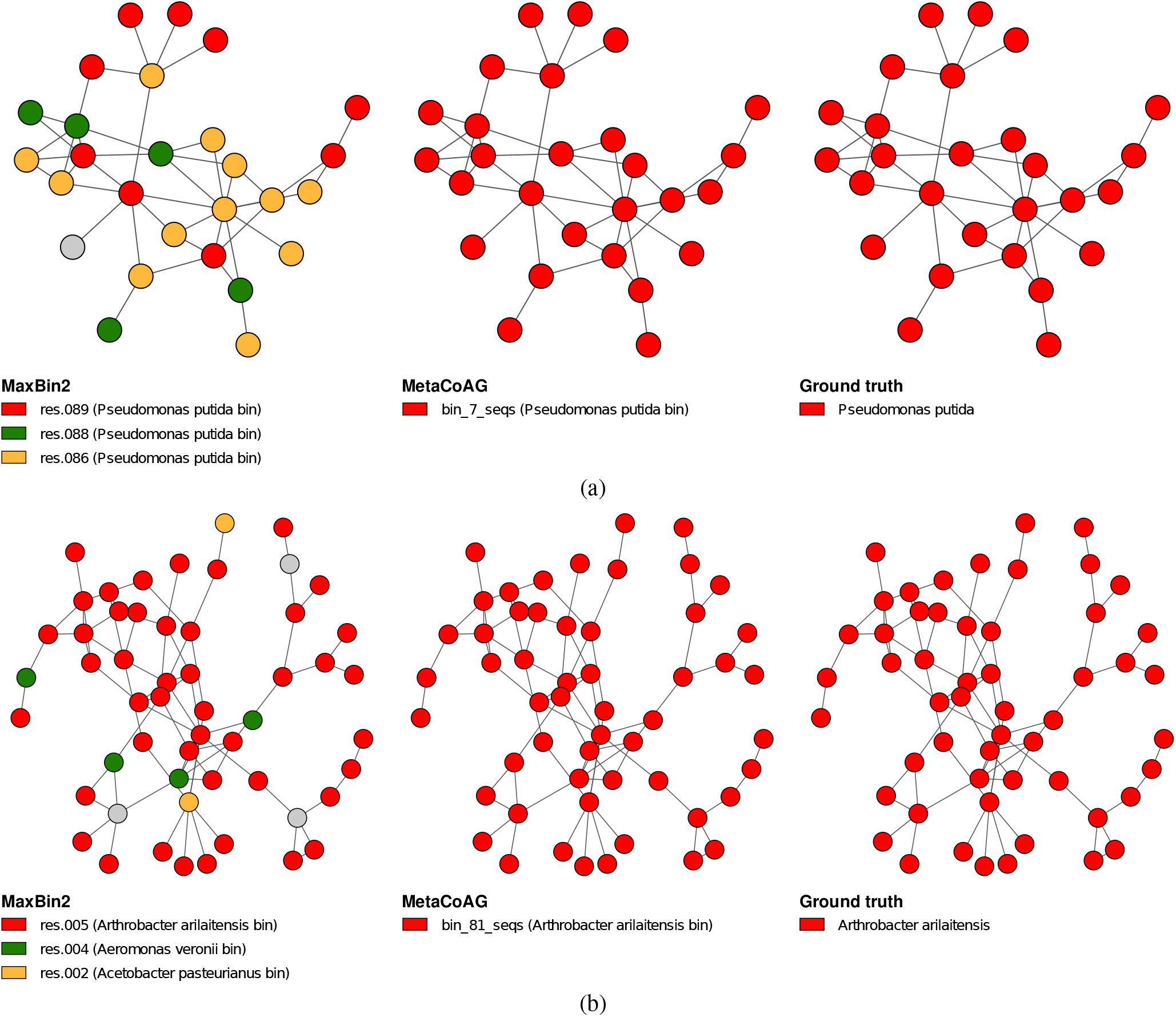
Visualization of the binning results of the simHC+ dataset for species with high variance in oligonucleotide composition and high variance in coverage. Visualization of the binning results from MaxBin2 and MetaCoAG for a species with (a) high variance in oligonucleotide composition (standard deviation > 0.015) and (b) high variance in coverage (standard deviation > 50×). Gray color nodes denote contigs which were binned to bins other than the ones specified in the figure.

Another challenge faced by the majority of the existing binning tools is the inability to accurately separate contigs of species belonging to the same genus, where such species tend to have similar oligonucleotide composition and appear in similar abundances. For example, the following three species in simHC+, *Streptococcus pneumoniae*, *Streptococcus thermophilus* and *Streptococcus suis* are in the same genus *Streptococcus*, and they have very similar oligonucleotide composition (Refer to Fig. 4 (a)) and similar coverages (*Streptococcus pneumoniae*: 56×, *Streptococcus thermophilus*: 60× and *Streptococcus suis*: 50×). Not surprisingly, contigs from these three species were incorrectly binned by MaxBin2 and even ignored by MataBAT2 because they share similar composition and coverage profiles (Refer to Fig. 4 (b)). On the contrary, MetaCoAG was able to accurately bin most of the contigs from these three species because they naturally form three subgraphs in the assembly graph (Refer to Fig. 4 (b)), thus improving the F1-scores of *Streptococcus pneumoniae* from 46.51% to 93.40%, *Streptococcus thermophilus* from 49.97% to 95.67% and *Streptococcus suis* from 72.39% to 95.95%. Fig. 4 (b) demonstrates that the use of assembly graph in MetaCoAG can assist in the separation of species, despite the high similarity in oligonucleotide composition and coverage of certain species.

**Figure 4.**
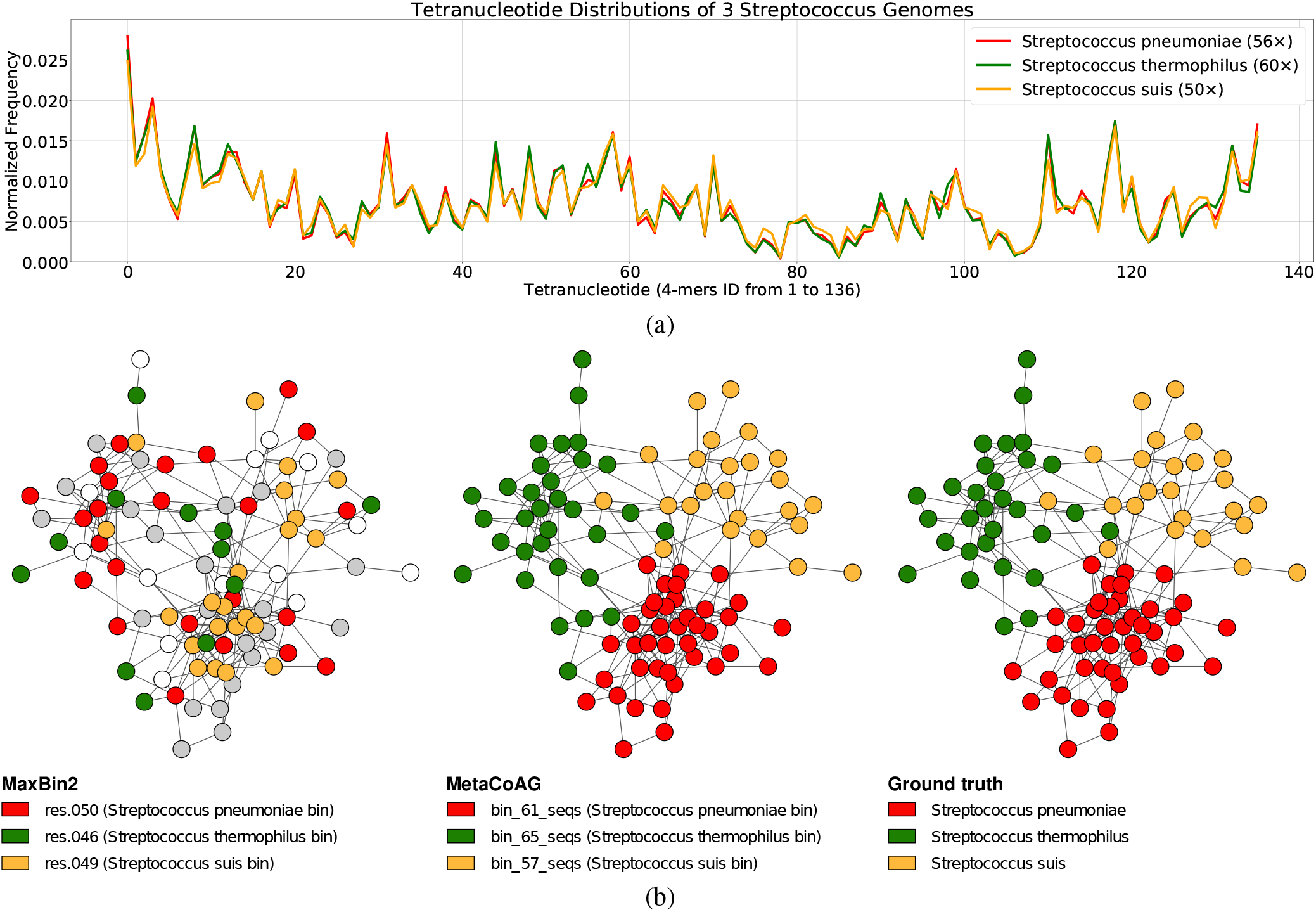
Visualization of the tetranucleotide composition and binning results of three *Streptococcus* genomes in the simHC+ dataset. (a) Tetranucleotide distributions of the three *Streptococcus* genomes; *Streptococcus pneumoniae* (red) with 56× coverage, *Streptococcus suis* (yellow) with 60× coverage and *Streptococcus thermophilus* (green) with 50× coverage. (b) Visualization of the binning results from MaxBin2 and MetaCoAG for three *Streptococcus* genomes. White color nodes denote discarded contigs and gray color nodes denote contigs which were binned to bins other than the three *Streptococcus* genomes. MetaBAT2 was not included as it had not recovered these three species.

### Benchmarks using CAMI2 Toy Human Microbiome Project Datasets

We benchmarked MetaCoAG against MaxBin2^18^, MetaBAT2^29^, and Vamb^30^ on five publicly available datasets from the toy Human Microbiome Project dataset of the second Critical Assessment of Metagenomic Interpretation (CAMI)^33^ challenge (Please refer to Supplementary Data 1 Table 1 for further details of the CAMI datasets). Multiple samples from each dataset were co-assembled together to obtain the final contigs for binning. Please refer to Supplementary Data 1 Fig. 5-7 for the multi-sample binning results, where we assembled the samples individually and binned them.

We evaluated the binning results of the CAMI datasets using CheckM^32^ and reported the F1-score of the bins produced by all the binning tools. Fig. 5 (a)–(e) shows that overall MetaCoAG has achieved the best binning results among all the binning tools. The overall median F1-scores averaging from all 5 CAMI datasets for MetaCoAG, MaxBin2, MetaBAT2 and Vamb are 86.77%, 75.41%, 1.57% and 33.30%, respectively. More specifically, MetaCoAG has recovered more complete bins with higher purity when compared to other tools (Please refer to Supplementary Data 1 Fig. 3 and 4 for completeness and purity results). MetaCoAG produced the highest numbers of high-quality and medium-quality bins combined together for all the CAMI datasets (Refer to Supplementary Data 1 Table 6). Note that only MaxBin2 outperforms MetaCoAG in terms of the number of high-quality bins just for the GI dataset. This dataset had a low density in its assembly graph (Please refer to Supplementary Data 1 Table 1 for density of the assembly graph) which prevented MetaCoAG from making full use of the assembly graphs.

**Figure 5.**
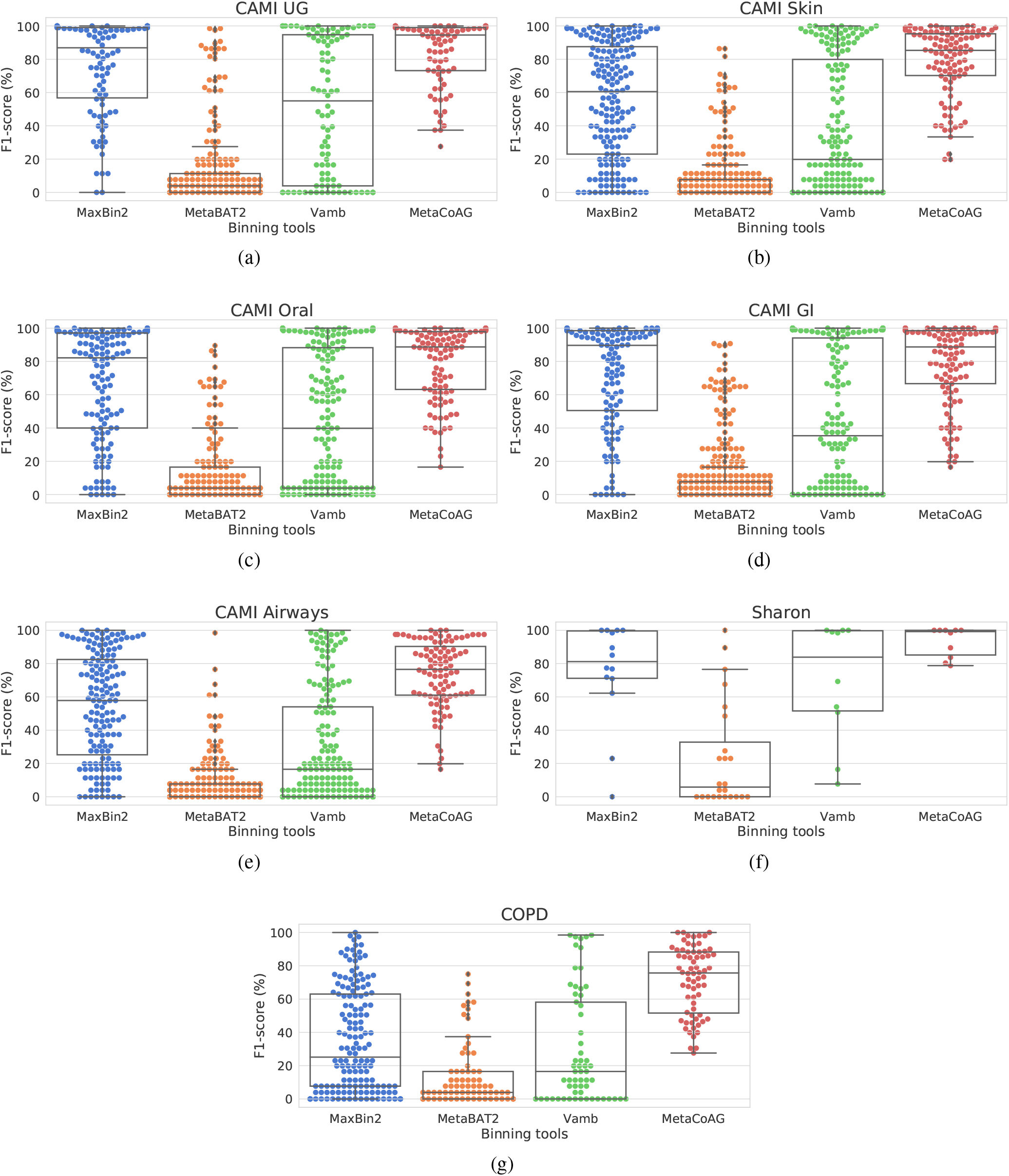
F1-score of the CAMI and real datasets. The F1-scores of the bins found in the CAMI and real datasets by all the binning tools in co-assembly mode.

### Benchmarks using Sharon and COPD datasets

We benchmarked MetaCoAG against MaxBin2^18^, MetaBAT2^29^ and Vamb^30^ on two real metagenomic datasets; **Sharon** dataset obtained from a pre-born infant’s gut^34^ and **COPD** dataset obtained from the Chronic Obstructive Pulmonary Disease (COPD) Lung Microbiome^35^. These datasets contain multiple samples (or runs) and further details about the samples used can be found in Supplementary Data 1 Table 3. The contigs and assembly graph for each dataset was obtained by co-assembling the reads from all the samples together. Please refer to Supplementary Data 1 Fig. 5-7 for the multi-sample binning results, where we assembled the samples individually and binned them.

Similar to the simHC+ and CAMI datasets, we again use CheckM^32^ to evaluate the bins produced by all the binning tools and identify high-quality bins. Fig. 5 (f)–(g) shows that MetaCoAG has also achieved the best binning result in terms of the median F1-score for both the real datasets. For the Sharon dataset, MetaCoAG records a median F1-score of 99.24% while the second-best tool (Vamb) has a median F1-score of 83.88%. For the COPD dataset, MetaCoAG records a median F1-score of 75.68% while the second-best tool (MaxBin2) has a median F1-score of 25.13%. Furthermore, MetaCoAG has produced the highest number of high-quality bins and the lowest number of low-quality bins for both the real datasets (Please refer to Supplementary Data 1 Table 6 for the exact counts).

We used GTDB-Tk^36^ to annotate all the high-quality bins produced by MetaCoAG, MaxBin2 and Vamb ^3^ for both datasets. Then we compared the taxonomic annotations (up to the species level) with the analysis results reported by the authors of these datasets (Refer to Table 1). Table 1 shows that MetaCoAG achieves the best consistency with the original analysis reported by the authors. In the Sharon dataset, the five most abundant species reported according to the authors^34^; *Staphylococcus epidermidis*, *Enterococcus faecalis*, *Cutibacterium avidum*, *Peptoniphilus lacydonensis* and *Staphylococcus aureus* have been successfully identified by all the three binning tools. However, Vamb missed *Staphylococcus hominis*, which is reported as a rare species in the Sharon dataset^34^. Moreover, MetaCoAG is the only tool that is able to recover *Leuconostoc citreum*, which is also identified as a rare species in the Sharon dataset^34^. These results denote the ability of MetaCoAG to recover rare species in real metagenomics samples that are ignored by other binning tools.

In the COPD dataset, there is a larger discrepancy among MaxBin2, Vamb and MetaCoAG. Only two species, *Peptostrepto-coccus sp.* and *SR1 bacterium human oral taxon HOT-345*, have been identified by all the three binning tools. *SR1 bacterium human oral taxon HOT-345* and *Lachnospiraceae bacterium oral taxon 096* have been added to NCBI taxonomy recently^37^ and hence are not found in the original analysis^35^. Compared to MetaCoAG, MaxBin2 failed to identify three species *Prevotella pallens*, *Prevotella shahii* and *Prevotella histicola* while Vamb only identified *Prevotella pallens* under the genus *Prevotella*. Similarly, Vamb failed to identify two species, *Capnocytophaga gingivalis* and *Capnocytophaga leadbetteri*, both of which are identified by MaxBin2 and MetaCoAG. Moreover, the species *Anaeroglobus micronuciformis* only identified by MaxBin2 was not present in the top 50 genera ranked by abundance in the original analysis^35^, which is likely to be a false-positive. These results demonstrate that MetaCoAG has been able to recover more species correctly with respect to the original analysis of these real datasets.

## Discussion

Metagenomic sequencing and *de novo* assembly, coupled with binning methods have facilitated the characterization of different microbial communities. The majority of existing metagenomic contig-binning tools do not make use of the valuable connectivity information found in assembly graphs from which the contigs are derived. Furthermore, existing tools do not make use of multiple single-copy marker genes throughout the entire binning process.

MetaCoAG is a stand-alone tool for binning metagenomic contigs that makes use of composition, coverage and assembly graphs simultaneously. The use of connectivity information from the assembly graphs makes the binning process of MetaCoAG robust against high variance of intra-species oligonucleotide composition and coverage as well as similar inter-species oligonucleotide composition and coverage (within the same genus). Experimental results on both simulated and real datasets show that MetaCoAG achieves the best binning results compared to state-of-the-art tools, especially producing more high-quality bins and recovering more species.

MetaCoAG can be easily extended to work with other assemblers based on assembly graphs (e.g., the de Bruijn graph^22^, the string graph^38^, the repeat graph^39^, etc.) for both short and long reads. In the future, we plan to extend MetaCoAG to support overlapped binning^25^, *i.e.* detect contigs that may belong to multiple species. Furthermore, we plan to incorporate MetaCoAG with assembly pipelines that may lead to more efficient and accurate analysis for metagenomic datasets.

## Methods

### Step 0: Assemble Reads into Contigs and Construct the Assembly Graph

This preprocessing step is carried out to assemble the reads into contigs and obtain the assembly graph. Metagenomic assemblers first use graph models to connect overlapping reads or *k*-mers and to infer contigs as non-branching paths. After graph simplification, the vertices represent contigs and edges represent connections between contigs in the assembly graph. Here we use the popular metagenomic assembler metaSPAdes^21^ to derive input contigs and assembly graphs. Note that the assembly graphs can also be obtained similarly using other metagenomic assemblers such as MEGAHIT^20^ and metaFlye^40^.

### Step 1: Identify Contigs with Single-Copy Marker Genes

Single-copy marker genes appear only once in a bacterial genome and are conserved in the majority of bacterial genomes^15–17^. Similar to approaches such as MaxBin^17^ and MaxBin2^18^, MetaCoAG uses 107 single-copy marker genes to distinguish contigs belonging to different species. For each of the 107 single-copy marker genes, we use FragGeneScan^41^ and HMMER^42^ to identify the contigs containing this single-copy marker gene. A single-copy marker gene is considered to be contained in a contig if at least 50% of the length of the gene is aligned to this contig.

### Step 2: Order Single-copy Marker Genes and Estimate the Number of Initial Bins

For a given single-copy marker gene, the contigs containing this marker gene should come from different species (*e.g.*, if two contigs contain the same marker gene, then the two contigs should belong to two different species). In the ideal case, if we have a near-perfect assembly, the number of contigs that contain the same single-copy marker gene should be equal to the number of species present in the sample. However, in reality, assemblies can be fragmented and erroneous, which may make it challenging to recover all single-copy marker genes and hence, lowering the counts of contigs containing each single-copy marker gene.

To get a better estimation of the number of species, we obtain the counts of contigs containing each single-copy marker gene. We also recorded the single-copy marker genes found in each contig. Now we order all the single-copy marker genes according to the descending order of the number of contigs containing them. For the single-copy marker genes having the same number of contigs, we order them according to the descending order of the total count of single-copy marker genes found in its constituent contigs. We refer to this list of ordered marker genes as *SMG* where a single-copy marker gene *g*_*i*_ has a set of contigs *C*(*g*_*i*_) containing *g*_*i*_. Then, the number of initial bins is set to be the largest count of contigs a marker gene has to recover the maximum number of species possible from the marker gene information.

### Step 3: Bin Contigs with Single-copy Marker Genes

#### Step 3a: Initialize Bins

We initialize the bins using the contigs of the first single-copy marker gene *g*_1_ in *SMG*; *i.e.*, we initialize a new bin *B* for each contig in *C*(*g*_1_) (as shown in Step 3a of Fig. 1). We define the initialized set of bins as *BINS*. Please note that the number of bins |*BINS*| may change during the binning process.

#### Calculating Composition and Coverage Probabilities

Previous studies on metagenomics binning have used genomic signatures as they follow species-specific patterns^17, 43^. The most commonly used genomic signatures to characterise composition information are *tetranucleotide frequencies* (strings of length *k* = 4, also known as *tetramers*). We obtain the normalized tetranucleotide frequency vectors of each contig *c* as *tetra*(*c*). We obtain the tetranucleotide composition distance *d*_*tetra*_(*c, c*′) between two contigs *c* and *c*′ as shown in equation 1 where *dist*_*E*_ is the Euclidean distance function.

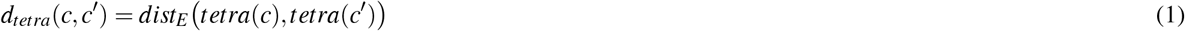

We follow the method used by Wu *et al.*^17^ and define the probability function that *c* and *c*′ belong to the same specie based on their composition, *P*_*comp*_(*c,c*′) as shown in equation 2.

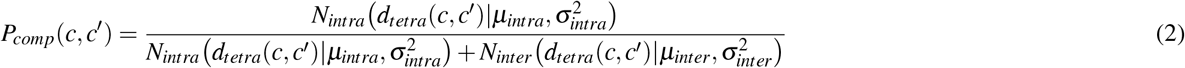

*N*_*intra*_ and *N*_*inter*_ are Gaussian distributions with *μ*_*intra*_, *σ*_*intra*_, *μ*_*inter*_ and *σ*_*inter*_ set according to the latest values of MaxBin 2.2.7^18^ which have been calculated by analysing the Euclidean distance between the tetranucleotide frequencies of pairs of sequences sampled from the same genome (*intra*) and different genomes (*inter*). If the distance is lower between two sequences, they are more probable to belong to the same genome.

We use the coverage information of the contigs as coverage carries important information about the abundance of species and has been used in previous metagenomics binning studies^15, 17^. Shotgun sequencing has shown to follow the Lader-Waterman model^44^ and the Poisson distribution has been used to obtain the sequencing coverage of nucleotides and applied in metagenomics binning^17, 45^. Modifying the definition found in Wu *et al.*^17^, we define the probability function that *c* and *c*′ belongs to the same species given their coverage values in each sample, *P*_*cov*_(*c, c*′) as shown in equation 3.

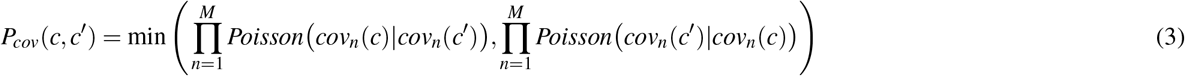

Here *cov*_*n*_(*c*) and *cov*_*n*_(*c*′) refer to the coverage values of the contigs *c* and *c*′ respectively in the sample *n* where *M* is the number of samples. *Poisson* is the Poisson probability mass function.

#### Step 3b: Construct a Weighted Bipartite Graph and Find a Minimum-Weight Full Matching

In the previous steps, we have used single-copy marker genes to identify pairs of contigs that belong to different species. Remind that contigs in different bins in *BINS* are expected to belong to different species and contigs in *C*(*g*_*i*_) are also expected to belong to different species. However, there is no measurement to measure how likely a contig *c* in *C*(*g*_*i*_) belongs to an existing bin *B* in *BINS*. Therefore, we introduce a bipartite graph between *C*(*g*_*i*_) and *BINS* and propose a weight *w*_*c2B*_(*c,B*) between a contig *c* in *C*(*g*_*i*_) and an existing bin *B* in *BINS* as shown in equation 4 (averaging over all the contigs in bin *B*).

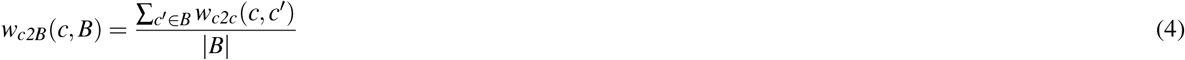

In equation 4, *w*_*c2c*_(*c,c*′) is the weight that measures how likely a pair of contigs *c* and *c*′ belong to the same species and is computed using equation 5.

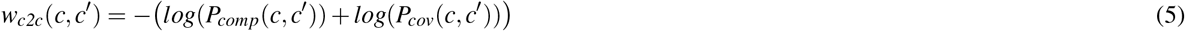

In equation 5, *P*_*comp*_(*c,c*′) and *P*_*cov*_(*c,c*′) are calculated according to equations 2 and 3 respectively.

Now we find a minimum-weight full matching (minimum-cost assignment)^46^ for the above bipartite graph between *C*(*g*_*i*_) and *BINS* where every contig *c* in *C*(*g*_*i*_) will get paired with exactly one bin *B* in *BINS*. For this purpose, we use the minimum-weight full matching algorithm implemented in the *NetworkX* ^4^ python library which is based on the algorithm proposed by Karp^46^ and the time complexity is *O*(|*C*(*g*_*i*_|) × |*BINS*| × *log*(|*BINS*|)).

In the next step, we will see how we can assign the contigs to existing bins based on the minimum-weight full matching we have obtained.

#### Step 3c: Assign Contigs to Existing Bins or Dynamically Adjust Bins

Previous studies have observed that contigs connected to each other in the assembly graph are more likely to belong to the same taxonomic group^23, 24^. While *w*_*c2B*_(*c,B*) considers both composition and coverage information, the assembly graph has not yet been incorporated into the binning process. Therefore, we introduce *d*_*graph*_(*c,B*) to measure how well contig *c* is connected to contigs in bin *B* within the assembly graph. Specifically, *d*_*graph*_(*c,B*) is defined as the average length of the shortest-path distances between contig *c* and all the contigs in bin *B* in the assembly graph. Note that both *w*_*c2B*_(*c,B*) and *d*_*graph*_(*c,B*) will be used to assign contigs to existing bins or dynamically adjust the bins.

We define the thresholds *w*_*intra*_ and *w*_*inter*_ as follows where *M* is the number of samples in the dataset.

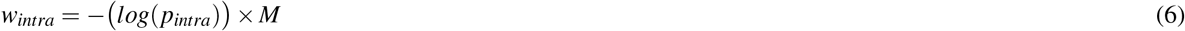

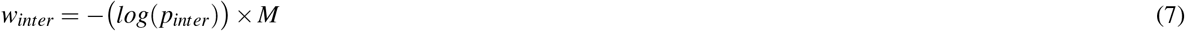

Each candidate pair (*c,B*) obtained from the minimum-weight full matching falls under one of the following three cases as shown in Supplementary Data 1 Fig. 1.

- **Case 1:** If the weight of the candidate pair *w*_*c2B*_(*c,B*) is less than or equal to *w*_*intra*_ and the average distance *d*_*graph*_(*c,B*) is less than or equal to *d*_*limit*_, then contig *c* will be assigned to bin *B*, *i.e.*, *B ← B ∪ {c}* (*e.g.*, contig 4 and Bin 1 in Supplementary Data 1 Fig. 1).
- **Case 2:** If the weight of the candidate pair *w*_*c2B*_(*c,B*) is greater than *w*_*inter*_ and the average distance *d*_*graph*_(*c,B*) is greater than *d_limit_*, then a new bin *B*′ is created and contig *c* is assigned to that new bin, *i.e.*, *B*′ = {*c*} and *BINS ← BINS ∪ {B}*′. (*e.g.*, contig 21 in Supplementary Data 1 Fig. 1).
- **Case 3:** If *w*_*c2B*_(*c,B*) and *d*_*graph*_(*c,B*) satisfy neither Case 1 nor Case 2, then contig *c* will not be assigned to any bin (*e.g.*, contig 14 in Supplementary Data 1 Fig. 1).

The default values for parameters *p*_*intra*_, *p*_*inter*_, *d*_*limit*_ were chosen empirically and set to 0.1, 0.01 and 20 respectively. Now we iteratively perform Steps 3b and 3c to process all the contigs containing single-copy marker genes. The remaining challenge is to bin the contigs which do not contain single-copy marker genes which will be addressed in Step 4.

### Step 4: Bin Remaining Contigs Using Label Propagation

After we bin the contigs with single-copy marker genes, each such contig receives a label corresponding to its bin. Now we will propagate labels from these contigs to other unlabeled contigs within the same connected component.

#### Step 4a: Propagate Labels Within Connected Components

MetaCoAG uses composition, coverage and distance information from the assembly graph to propagate labels from labeled contigs to the unlabeled contigs located within the same connected components. More specifically, for each unlabeled long contig *c* (at least 1,000 bp long because short contigs result in unreliable composition and coverage information) directly connected or connected via short contigs to a labeled contig *c*′, MetaCoAG computes a candidate propagation action (*c*′,*c,d*(*c,c*′), *w*_*c2B*_(*c,B*′)) where *d*(*c,c*′) is the shortest distance between *c* and *c*′ using only unlabeled vertices and *w*_*c2B*_(*c, B*′) is computed according to equation 4 where *B*′ is the bin to which contig *c*′ is assigned. Given two candidate propagation actions (*a,b,d,w*) and (*a*′,*b*′,*d*′,*w*′), (*a,b,d,w*) has a higher priority than (*a*′, *b*′, *d*′, *w*′) if *d* <*d*′ or (*w* < *w*′ and *d* = *d*′). MetaCoAG iteratively selects the candidate propagation action with the highest priority and executes the corresponding label propagation. If a contig to be labeled contains single-copy marker genes, the relevant candidate propagation action is executed if the single-copy marker genes of the contig are not present in the intended bin. We restrict the depth of the search for labeled contigs in this step to 10 in order to speed up MetaCoAG.

#### Step 4b: Propagate Labels Across Different Components

Note that some components in the assembly graph may not have any labeled contigs and we need to propagate labels from labeled bins to unlabeled contigs across components. Calculating pair-wise weights *w*_*c2c*_(*c,c*′) for all the remaining contigs becomes time consuming. Hence, for each bin *B* we create a representative contig *c*(*B*) which has a composition profile and a coverage profile calculated by averaging the normalized tetranucleotide frequency vectors and coverage vectors of all the contigs in bin *B*, respectively. These profiles will provide a better representation of the composition and coverage of the bins. Then, for each unlabeled contig *c*, MetaCoAG identifies a bin *B* that minimizes *w*_*c2c*_(*c, c*(*B*)) which is calculated according to equation 5, and assigns contig *c* into that bin *B*. This propagation is limited to long contigs (at least 1,000 bp long by default). If an unlabeled contig contains single-copy marker genes, it is assigned to bin *B* that minimizes *w*_*c2c*_(*c,c*(*B*)) if the single-copy marker genes of the contig are not present in bin *B*. Then, Step 4a is performed again to further propagate labels.

#### Step 4e: Postprocessing

In this step, we will make final adjustment on the current bins. Two bins *B* and *B*′ are *mergeable* if they have no common marker genes and *w*_*c2c*_(*c*(*B*), *c*(*B*′)) (calculated by equation 5) is upper bounded by *w*_*intra*_ (defined in Step 3c). Then, MetaCoAG creates a graph where vertices denote current bins and edges between two vertices denote that the corresponding two bins are mergeable. Now we use the implementation of python-igraph library to find maximal cliques (https://igraph.org/c/doc/igraph-Cliques.html#igraph_maximal_cliques) in this graph and merge the bins found in each maximal clique. After merging bins, we also remove the bins which contain less than one third (set by default) of the single-copy marker genes. Finally, MetaCoAG outputs the bins along with their corresponding contigs.

### Datasets

#### Simulated Datasets

We evaluated the binning performance on the simulated **simHC+** dataset^17^ which consists of 100 bacterial species. Paired-end MiSeq reads were simulated using InSilicoSeq^47^ with 300 bp mean read length.

#### CAMI2 Toy Human Microbiome Project Datasets

We used the simulated metagenome data from the toy Human Microbiome project of the second CAMI challenge^33^. Metagenomes were simulated from five different body sites of the human host as follows.

1. Urogenital tract - referred as **CAMI UG**
2. Skin - referred as **CAMI Skin**
3. Oral cavity - referred as **CAMI Oral**
4. Gastrointestinal tract - referred as **CAMI GI**
5. Airways - referred as **CAMI Airways**

#### Real Datasets

We used the following real datasets to evaluate the binning performance on real-world metagenomic data.

1. Pre-born infant gut metagenome,^34^ - referred as **Sharon**
2. Metagenomics of the Chronic Obstructive Pulmonary Disease (COPD) Lung Microbiome^35^ - referred as **COPD**

Please refer to Supplementary Data 1 Tables 1 and 3 for further details of all the datasets.

### Tools Used

We used the popular metagenomic assembler metaSPAdes^21^ (from SPAdes version 3.15.2^48^) to assemble reads into contigs and obtain the assembly graphs. The mean coverage of each contig in each sample was calculate using CoverM (available at https://github.com/wwood/CoverM).

MetaCoAG was benchmarked against the binning tools MaxBin2 (version 2.2.7) ^18^ in its default settings, MetaBAT2 (version 2.12.1)^29^ with `-m 1500` and Vamb (version 3.0.1)^30^ in both co-assembly and multi-sample modes with the parameter `--minfasta 200000` as suggested by the authors. The commands used to run these tools can be found in Supplementary Data 1 Fig. 7.

The binning results were evaluated using the tools AMBER^31^ (version 2.0.2), CheckM^32^ (version 1.1.3) and GTDB-Tk^36^ (version 1.5.0).

### Evaluation Criteria

Since the ground truth species for the simHC+ dataset were available, we used Minimap2^49^ to map the contigs to the reference genomes and determine the ground truth. With this ground truth annotation of contigs, we used AMBER^31^ to assess the binning results of the simHC+ dataset. We set the recall as AMBER completeness and precision as AMBER purity and calculated the F1-score as 2 ×(precision×recall)/(precision+recall) for each bin/species.

For all the datasets, we determined the completeness and contamination of the bins produced by each tool using CheckM^32^. We set the completeness as CheckM completeness and purity as 1/(1 + CheckM contamination). To check the trade-off between completeness and purity, we set the recall as completeness and precision as purity, and calculated the F1-score as 2 ×(precision×recall)/(precision+recall) for each bin. Furthermore, we counted the number of high-quality bins (bins which have >80% recall and >90% precision), medium-quality bins (bins which have >50% recall and >80% precision) and low-quality bins (bins which are not considered as high-quality or medium-quality).

To determine the species identified by the binning tools, we annotated all the high-quality bins of the real metagenomic datasets produced from the three best-performing tools; MetaCoAG, MaxBin2 and Vamb using GTDB-Tk^36^ up to the species level. The species were determined by the classification string produced by GTDB-Tk.

## Data availability

All the CAMI and real datasets containing raw sequencing data used for this study are publicly available from their respective studies. The CAMI2 Toy Human Microbiome Project datasets were downloaded from https://data.cami-challenge.org/participate from the 2nd CAMI Toy Human Microbiome Project Dataset. The Sharon dataset was downloaded from NCBI with BioProject number PRJNA60717 and accession number SRA052203. The COPD dataset was downloaded from NCBI with BioProject number PRJEB9034. NCBI accession numbers of the runs used to assemble the Sharon and COPD datasets can be found in Supplementary Data 1 Table 3. All the assembled data and results from all the binning tools, including the source data for Figs. 2–5 and Table 1 are available on figshare at https://figshare.com/projects/MetaCoAG/121014.

## Code availability

The code of MetaCoAG can be found on GitHub at https://github.com/Vini2/MetaCoAG and is freely available under the GPL-3.0 license. All analyses in this paper were performed using MetaCoAG v.1.0 with default parameters.

## Acknowledgements

This research was undertaken with the assistance of resources and services from the National Computational Infrastructure (NCI), which is supported by the Australian Government.

## Author contributions

V.M. and Y.L. designed the algorithm and drafted the manuscript. V.M. implemented MetaCoAG and conducted the experiments. V.M. and Y.L. analyzed the results. Y.L. supervised the project. All authors reviewed the manuscript.

## Competing interests

The authors declare that they have no competing interests.

## Footnotes

1 Please note that the recently published tool Vamb30 was not used to evaluate the simHC+ dataset as the number of contigs was less than the number recommended by the authors (https://github.com/RasmussenLab/vamb#recommended-workflow).

2 MetaBAT2 was not included in this comparison as it had not recovered the species *Pseudomonas putida* and *Arthrobacter arilaitensis*.

3 MetaBAT2 results were not considered for GTDB-Tk annotations as the results had very low number of high-quality bins compared to the other binning tools.

4 https://networkx.org/documentation/stable/reference/algorithms/generated/networkx.algorithms.bipartite.matching.minimum_weight_full_matching.html

## Notes

### Competing Interest Statement

The authors have declared no competing interest.

