## Supplementary Data 1 for "MetaCoAG: Binning Metagenomic Contigs via Composition, Coverage and Assembly Graphs"

### Supplementary Material for "MetaCoAG: Binning Metagenomic Contigs via Composition, Coverage and Assembly Graphs"

Vijini Mallawaarachchi and Yu Lin
School of Computing, Australian National University, Canberra, ACT 2600, Australia

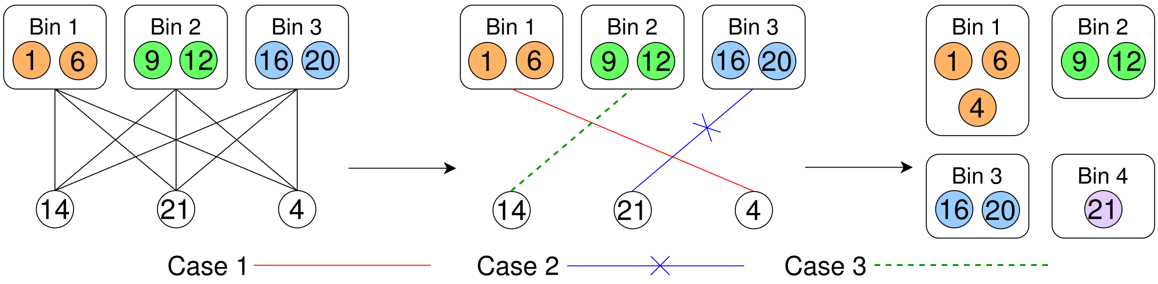


Figure 1: Cases 1, 2 and 3 in assigning contigs to existing bins or adjusting bins.

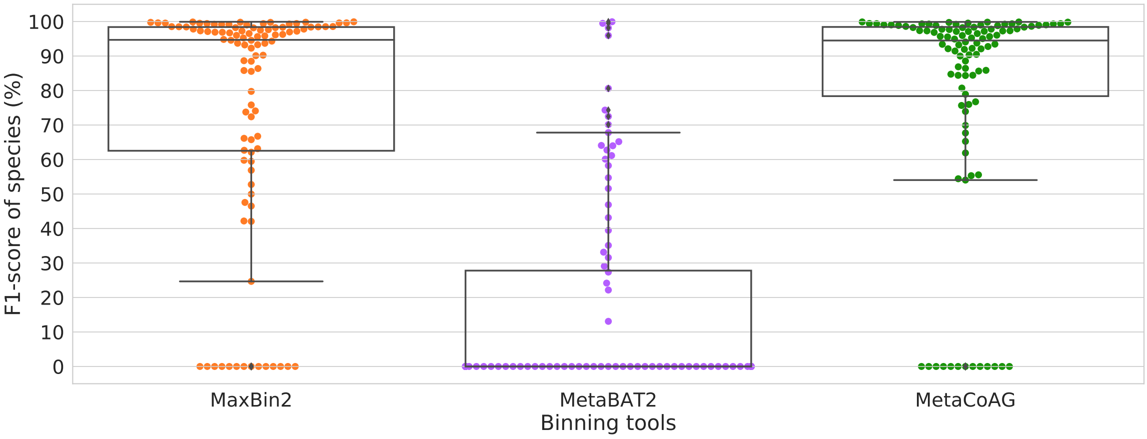


Figure 2: F1-score of species recovered by all the binning tools for the simHC+ dataset.

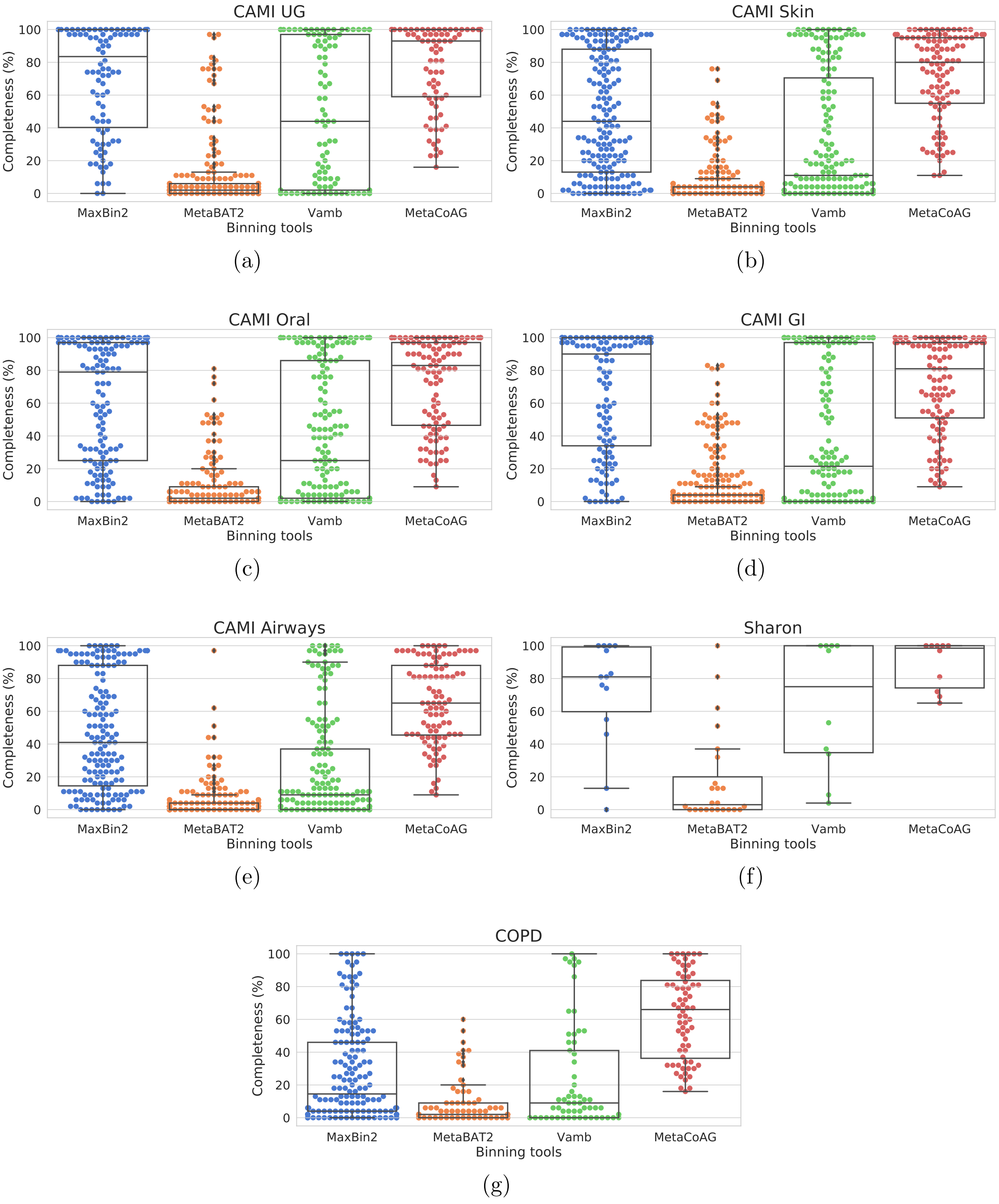


Figure 3: Completeness results of the CAMI and real datasets by all the binning tools in co-assembly mode.

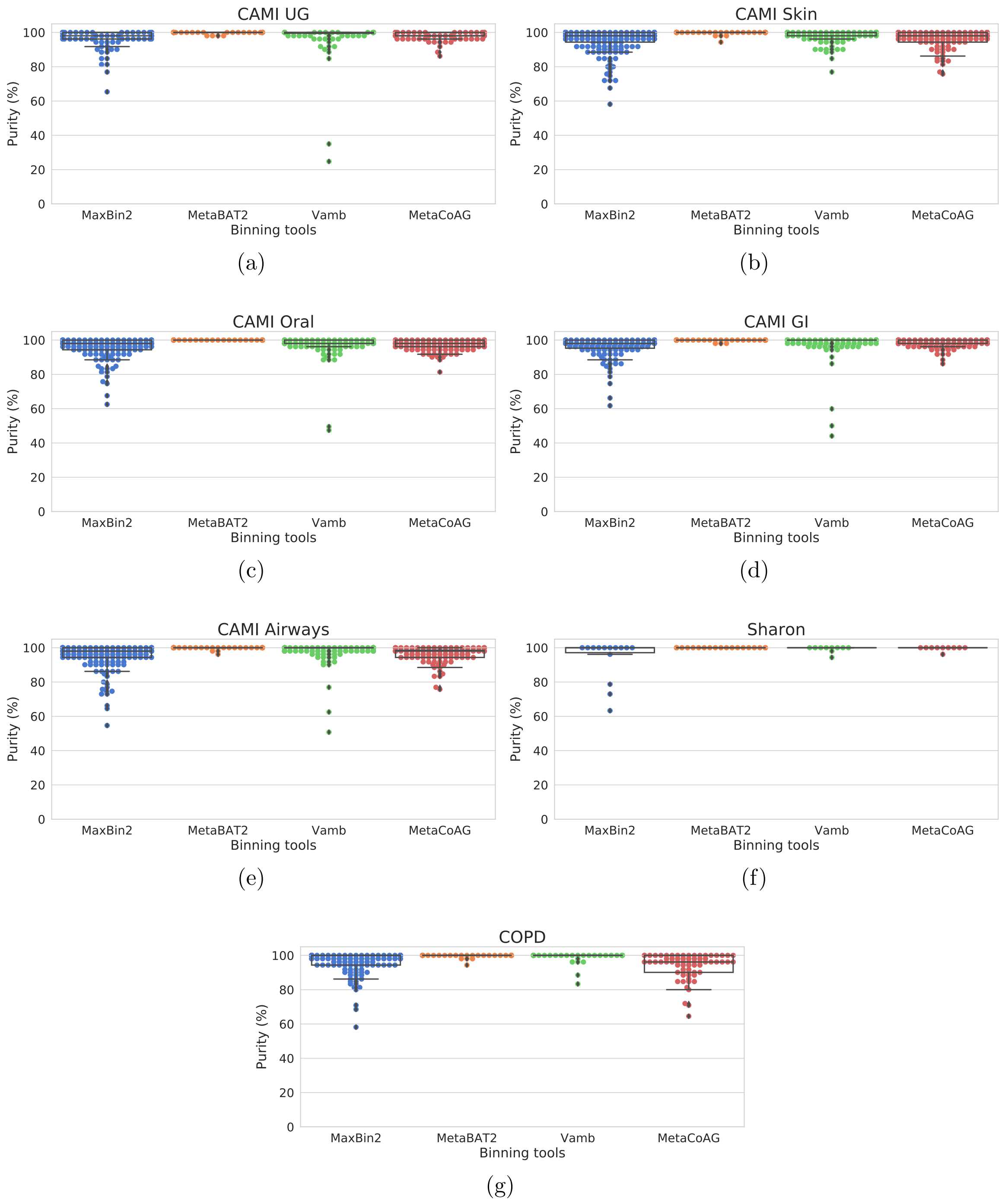


Figure 4: Purity results of the CAMI and real datasets by all the binning tools in co-assembly mode.

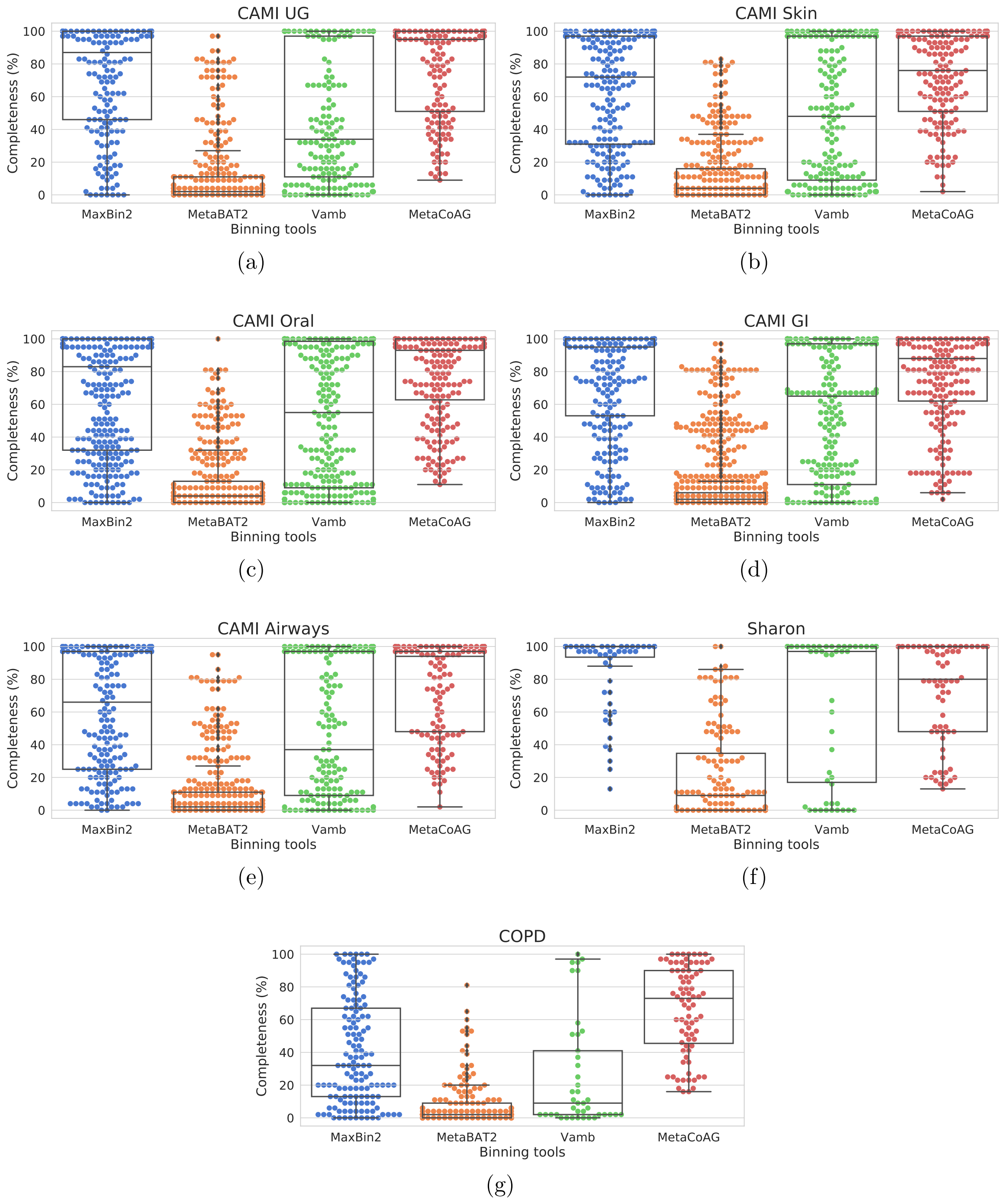


Figure 5: Completeness results of the CAMI and real datasets by all the binning tools in multi-sample mode.

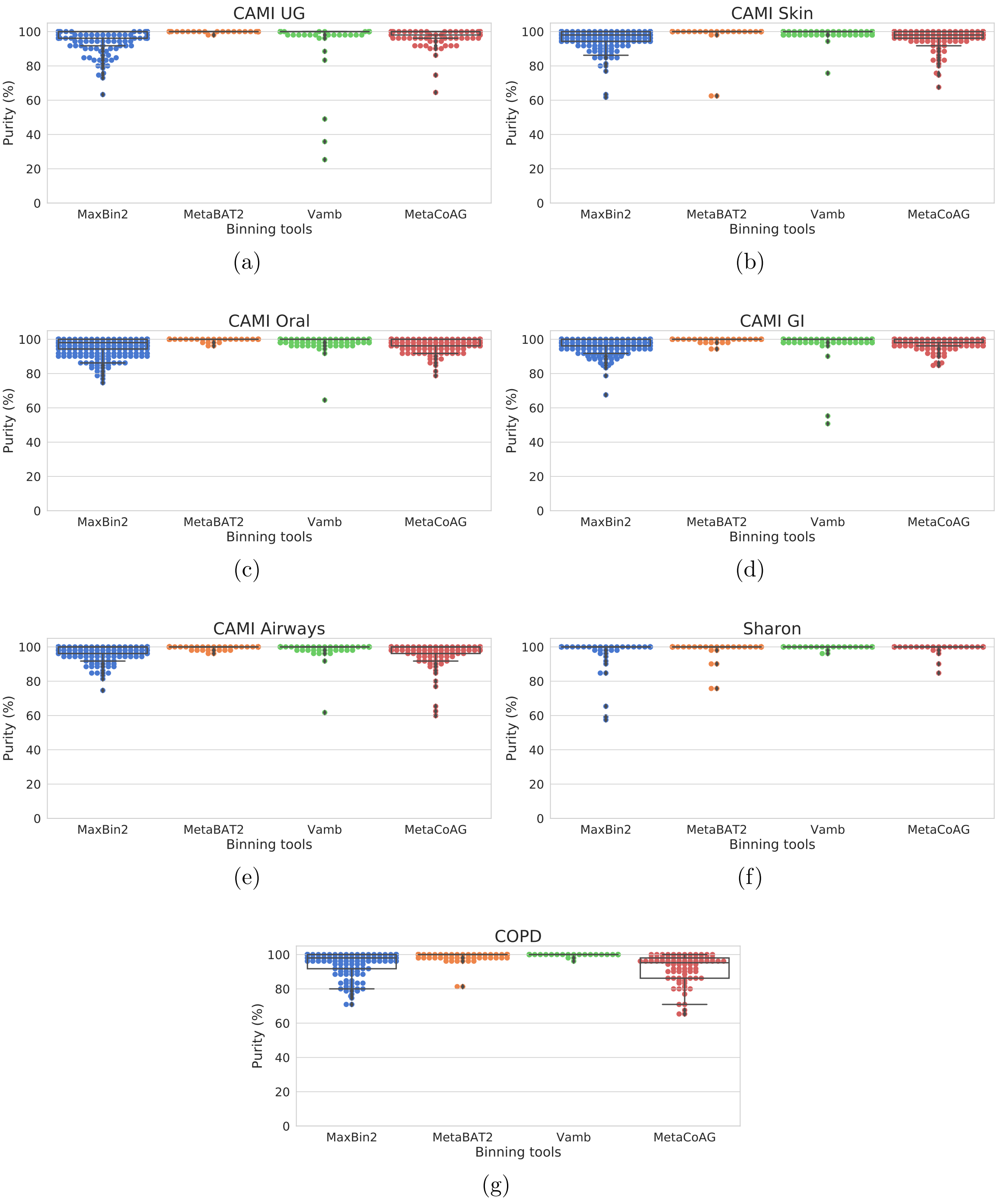


Figure 6: Purity results of the CAMI and real datasets by all the binning tools in multi-sample mode.

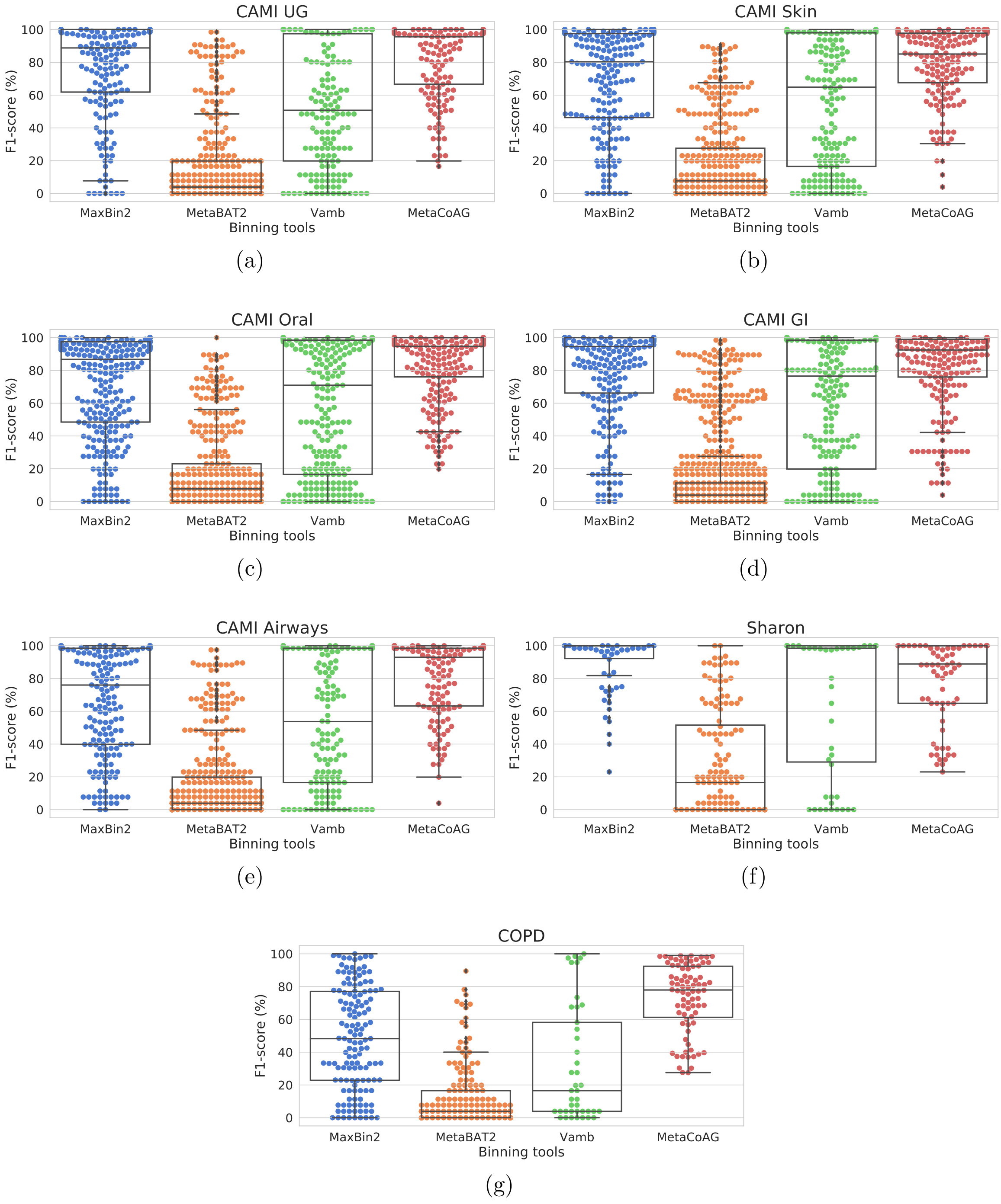


Figure 7: F1-score results of the CAMI and real datasets by all the binning tools in multi-sample mode.

**metaSPAdes**
```
> spades --meta -1 Reads_1.fastq -2 Reads_2.fastq -o /path/output_folder -t 32
```

**MaxBin2**
```
> perl MaxBin-2.2.7/run_MaxBin.pl -contig contigs.fasta -abund abundance.abund
-thread 8 -out /path/output_folder
```

**MetaBAT2**
```
> jgi_summarize_bam_contig_depths --outputDepth depth.txt *.bam
> metabat2 -i contigs.fasta -a depth.txt -o /path/output_folder/bin
```

**Vamb**
```
> minimap2 -d catalogue.mmi contigs.fasta
> minimap2 -t 8 -N 50 -ax sr catalogue.mmi R1.fastq R2.fastq | samtools view
-F 3584 -b --threads 8 > reads.bam
> vamb --outdir /path/output_folder --fasta contigs.fasta --bamfiles *.bam
--minfasta 200000
```

**MetaCoAG**
```
> ./MetaCoAG --assembler spades --contigs contigs.fasta --graph
assembly_graph_with_scaffolds.gfa --paths contigs.paths --abundance
abundance.tsv --output /path/output_folder
```

Figure 8: Commands used to run different tools.

| Dataset | No. of samples | Read length (bp) | Assembly size (Mb) | Total no. of assembled contigs | No. of contigs longer than 1000 bp | No. of edges in the assembly graph | Density† of the assembly graph | N50 (bp) |
|---|---|---|---|---|---|---|---|---|
| simHC+ [7] | 1 | 301 | 314.23 | 15,729 | 6,706 | 31,199 | 1.984 | 154,030 |
| CAMI UG [4] | 9 | 150 | 336.69 | 192,679 | 49,927 | 40,861 | 0.212 | 8,081 |
| CAMI Skin [4] | 10 | 150 | 639.91 | 600,508 | 139,217 | 51,139 | 0.085 | 1,965 |
| CAMI Oral [4] | 10 | 150 | 517.35 | 493,149 | 96,079 | 87,910 | 0.178 | 2,405 |
| CAMI GI [4] | 10 | 150 | 507.09 | 255,722 | 75,913 | 39,240 | 0.153 | 9,319 |
| CAMI Airways [4] | 10 | 150 | 664.49 | 729,063 | 142,477 | 49,531 | 0.067 | 1,506 |
| Sharon [5] | 18 | 100 | 45.06 | 37,164 | 7,067 | 20,328 | 0.547 | 5,609 |
| COPD [1] | 18 | 151 | 343.52 | 452,600 | 67,753 | 73,519 | 0.162 | 1,042 |

Table 1: Further information of the co-assemblies of the datasets. †Density of the graph is calculated as (the number of edges) / (the number of nodes).

| Evaluation criteria | MaxBin | MetaBAT2 | MetaCoAG |
|---|---|---|---|
| Average purity per bin (AMBER) | 90.36 | **98.30** | 91.07 |
| Average purity per bin (CheckM) | 97.25 | **100.0** | 97.55 |
| Average completeness per bin (AMBER) | 79.34 | 13.02 | **82.73** |
| Average completeness per bin (CheckM) | 77.51 | 29.59 | **87.17** |
| F1-score per bin (AMBER) | 84.50 | 23.00 | **86.70** |
| F1-score per bin (CheckM) | 80.64 | 37.25 | **89.44** |
| Accuracy | 77.07 | 14.38 | **84.46** |
| Binned fraction | 84.90 | 14.79 | **92.04** |

Table 2: AMBER and CheckM results for the simHC+ dataset.

| Dataset | BioProject number | NCBI accession numbers of the samples (runs) used |
|---|---|---|
| Sharon [5] | PRJNA60717 | SRR492197, SRR492196, SRR492195, SRR492194, SRR492193, SRR492192, SRR492191, SRR492190, SRR492189, SRR492188, SRR492187, SRR492186, SRR492185, SRR492184, SRR492183, SRR492182, SRR492066, SRR492065 |
| COPD [1] | PRJEB9034 | ERR970477, ERR970476, ERR970475, ERR970474, ERR970473, ERR970472, ERR970471, ERR970470, ERR970407, ERR970406, ERR970405, ERR970404, ERR970403, ERR970402, ERR970401, ERR970400, ERR970399, ERR970398 |

Table 3: Information of the samples used for the real datasets.

| Species | NCBI accession number | Relative Abaundance (%) | Coverage (×) | MaxBin2 F1-score (%) | MetaBAT2 F1-score (%) | MetaCoAG F1-score (%) |
|---|---|---|---|---|---|---|
| Acetobacter pasteurianus | AP011170.1 | 2.30 | 1533 | 96.92 | - | 98.85 |
| Aeromonas veronii | NC_015424.1 | 1.90 | 979 | 96.52 | 22.16 | 97.18 |
| Amycolatopsis mediterranei | CP003729.1 | 1.30 | 348 | 97.21 | 63.99 | 97.33 |
| Arthrobacter arilaitensis | NC_014550.1 | 1.20 | 557 | 97.65 | - | 98.99 |
| Azorhizobium caulinodans | NC_009937.1 | 1.20 | 269 | 85.55 | 67.81 | 85.65 |
| Bacillus cereus | NC_011658.1 | 1.10 | 251 | 94.33 | - | 88.56 |
| Bacillus thuringiensis | NC_014171.1 | 1.10 | 69 | 47.56 | - | - |
| Bdellovibrio bacteriovorus | NC_005363.1 | 1.10 | 49 | 99.81 | 99.95 | 99.83 |
| Bifidobacterium adolescentis | NC_008618.1 | 1.10 | 88 | 98.40 | - | 96.52 |
| Bifidobacterium animalis | CP001892.1 | 1.10 | 95 | 94.58 | 46.89 | 92.23 |
| Brachyspira intermedia | CP002874.1 | 1.10 | 46 | 99.41 | - | 99.90 |
| Campylobacter jejuni | NC_002163.1 | 1.10 | 94 | 99.88 | - | 99.88 |
| Candidatus Pelagibacter ubique | NC_007205.1 | 1.10 | 117 | 99.17 | - | 99.31 |
| Candidatus Phytoplasma mali | NC_011047.1 | 1.10 | 255 | 93.69 | - | 95.25 |
| Candidatus Sulcia muelleri | NC_013123.1 | 1.10 | 554 | 88.65 | - | 91.46 |
| Chlamydia trachomatis | CP002054.1 | 1.10 | 147 | 95.25 | - | 97.68 |
| Chlamydophila psittaci | CP002807.1 | 1.10 | 131 | 98.49 | 74.33 | 84.43 |
| Clostridium acetobutylicum | CP002118.1 | 1.10 | 39 | - | 64.09 | 65.26 |
| Clostridium botulinum | NC_015425.1 | 1.10 | 55 | - | - | 84.74 |
| Clostridium tetani | NC_004557.1 | 1.10 | 55 | - | - | 90.00 |
| Clostridium thermocellum | NC_009012.1 | 1.10 | 40 | - | - | 96.04 |
| Corynebacterium diphtheriae | NC_016788.1 | 1.10 | 62 | 65.76 | 43.14 | 55.56 |
| Corynebacterium pseudotuberculosis | NC_017462.1 | 1.10 | 66 | 92.29 | - | 93.23 |
| Corynebacterium ulcerans | CP002790.1 | 1.10 | 61 | 86.39 | - | 54.43 |
| Cyanobacterium UCYN | NC_013771.1 | 1.10 | 106 | 96.72 | 61.16 | 85.86 |
| Cyanothece sp | NC_014501.1 | 1.10 | 25 | 99.77 | - | 92.77 |
| Desulfovibrio vulgaris | NC_002937.3 | 1.10 | 25 | 99.81 | 33.14 | 99.75 |
| Ehrlichia ruminantium | NC_006831.1 | 1.10 | 102 | 96.90 | - | 98.38 |
| Enterococcus faecium | CP003351.1 | 1.10 | 52 | 95.85 | - | 97.81 |
| Erysipelothrix rhusiopathiae | NC_015601.1 | 1.00 | 86 | 98.17 | - | 86.43 |
| Escherichia coli | NC_011415.1 | 1.00 | 31 | - | - | - |
| Fervidicoccus fontis | NC_017461.1 | 1.00 | 116 | 99.63 | 80.62 | - |
| Fibrobacter succinogenes | CP002158.1 | 1.00 | 40 | 99.93 | 51.60 | 75.97 |
| Flavobacterium branchiophilum | NC_016001.1 | 1.00 | 43 | 98.24 | - | 99.20 |
| Francisella novicida | NC_008601.1 | 1.00 | 80 | 62.68 | - | 55.32 |
| Francisella tularensis | NC_009749.1 | 1.00 | 81 | 24.67 | - | - |
| Fusobacterium nucleatum | NC_003454.1 | 1.00 | 71 | 98.24 | - | 99.05 |
| Gardnerella vaginalis | CP002725.1 | 1.00 | 89 | 96.98 | 54.71 | 98.36 |
| Granulicella tundricola | NC_015064.1 | 1.00 | 36 | 98.52 | 24.17 | 98.59 |
| Haemophilus influenzae | NC_009567.1 | 1.00 | 81 | - | - | 97.37 |
| Haemophilus parainfluenzae | FQ312002.1 | 1.00 | 74 | - | - | 95.05 |
| Haemophilus somnus | NC_010519.1 | 1.00 | 68 | - | - | 97.87 |
| Halobacterium sp | AE004437.1 | 1.00 | 76 | 98.47 | - | - |
| Halothiobacillus neapolitanus | NC_013422.1 | 1.00 | 59 | 99.51 | - | 99.36 |
| Helicobacter pylori | CP001680.1 | 1.00 | 98 | 99.78 | - | 99.82 |
| Hyphomicrobium sp | NC_015717.1 | 1.00 | 32 | 90.25 | 72.49 | 90.49 |
| Ignavibacterium album | NC_017464.1 | 1.00 | 42 | 99.43 | 27.36 | 99.04 |
| Klebsiella oxytoca | NC_016612.1 | 1.00 | 26 | 52.76 | - | 80.72 |
| Krokinobacter sp | NC_015496.1 | 1.00 | 45 | 99.43 | 60.11 | 99.45 |
| *Continued to next page ...* | | | | | | |

Table 4: Information about the recovered species for the simHC+ dataset. "-" represents that the species was not recovered by the binning tool.

| Species | NCBI accession number | Relative Abaundance (%) | Coverage (×) | MaxBin2 F1-score (%) | MetaBAT2 F1-score (%) | MetaCoAG F1-score (%) |
|---|---|---|---|---|---|---|
| Lactobacillus brevis | NC_008497.1 | 1.00 | 67 | 79.75 | - | 98.26 |
| Lactobacillus casei | CP002618.1 | 1.00 | 50 | 93.29 | - | 98.91 |
| Lactobacillus delbrueckii | NC_008054.1 | 1.00 | 82 | 97.31 | - | 97.27 |
| Lawsonia intracellularis | NC_008011.1 | 1.00 | 105 | 75.83 | 98.17 | 73.93 |
| Legionella pneumophila | NC_014125.1 | 1.00 | 44 | 96.27 | 70.13 | 95.55 |
| Metallosphaera cuprina | NC_015435.1 | 0.90 | 83 | 99.63 | 65.17 | - |
| Methanocorpusculum labreanum | NC_008942.1 | 0.90 | 85 | 99.61 | - | - |
| Methanosarcina acetivorans | AE010299.1 | 0.90 | 27 | 74.10 | - | - |
| Methanosarcina barkeri | NC_007355.1 | 0.90 | 32 | 56.89 | - | - |
| Micrococcus luteus | NC_012803.1 | 0.90 | 61 | 85.81 | - | 84.35 |
| Mycobacterium bovis | CP002095.1 | 0.90 | 35 | - | - | 96.85 |
| Mycobacterium sp | NC_008146.1 | 0.90 | 27 | 42.07 | - | 86.88 |
| Mycoplasma gallisepticum | CP003512.1 | 0.90 | 157 | 97.28 | - | 97.77 |
| Mycoplasma hyorhinis | CP002669.1 | 0.90 | 185 | 97.81 | - | 99.39 |
| Neisseria meningitidis | CP001561.1 | 0.90 | 68 | 97.51 | - | 97.13 |
| Nitrosococcus watsonii | NC_014315.1 | 0.90 | 46 | 99.56 | 29.08 | 99.33 |
| Nitrosomonas sp | NC_015222.1 | 0.90 | 48 | 93.71 | - | 92.12 |
| Nocardia farcinica | NC_006361.1 | 0.90 | 26 | 42.18 | 31.56 | 69.90 |
| Odoribacter splanchnicus | NC_015160.1 | 0.90 | 35 | 96.13 | - | 99.05 |
| Paenibacillus mucilaginosus | NC_017672.1 | 0.90 | 14 | 88.49 | 13.08 | 93.45 |
| Paenibacillus sp | NC_013406.1 | 0.90 | 17 | 96.00 | 58.24 | 93.77 |
| Photobacterium profundum | NC_006370.1 | 0.90 | 30 | 94.56 | - | 95.72 |
| Prochlorococcus marinus | NC_009091.1 | 0.90 | 75 | 94.81 | - | 98.81 |
| Pseudogulbenkiania sp | NC_016002.1 | 0.90 | 28 | 97.82 | - | 98.23 |
| Pseudomonas putida | CP002290.1 | 0.90 | 21 | 59.78 | - | 99.56 |
| Rhizobium leguminosarum | NC_008380.1 | 0.90 | 24 | 93.22 | 35.11 | 94.09 |
| Rhodococcus jostii | NC_008268.1 | 0.90 | 16 | - | - | 92.09 |
| Rickettsia prowazekii | CP003391.1 | 0.90 | 111 | 95.64 | - | 94.92 |
| Rickettsia rickettsii | NC_016909.1 | 0.90 | 97 | - | - | - |
| Rickettsia slovaca | NC_016639.1 | 0.90 | 96 | - | - | 67.69 |
| Ruegeria sp | NC_008044.1 | 0.90 | 38 | 97.11 | 62.75 | 97.11 |
| Salmonella enterica | NC_011083.1 | 0.90 | 25 | 66.15 | - | 76.74 |
| Sebaldella termitidis | NC_013517.1 | 0.90 | 28 | 99.16 | 39.46 | 99.40 |
| Shewanella sp | NC_008322.1 | 0.90 | 26 | 99.21 | - | 99.25 |
| Shigella flexneri | NC_004741.1 | 0.90 | 27 | 73.77 | - | 78.92 |
| Sodalis glossinidius | NC_007712.1 | 0.90 | 29 | - | - | 75.67 |
| Staphylococcus aureus | CP001844.2 | 0.90 | 44 | 99.29 | - | 99.29 |
| Streptococcus pneumoniae | NC_010582.1 | 0.90 | 56 | 46.51 | - | 93.40 |
| Streptococcus pyogenes | NC_004606.1 | 0.90 | 65 | 98.55 | - | 97.30 |
| Streptococcus suis | CP002570.1 | 0.90 | 60 | 72.39 | - | 95.95 |
| Streptococcus thermophilus | NC_008532.1 | 0.90 | 50 | 49.97 | - | 95.67 |
| Streptomyces scabiei | NC_013929.1 | 0.90 | 9 | 66.74 | - | 84.41 |
| Symbiobacterium thermophilum | NC_006177.1 | 0.90 | 26 | 98.36 | - | 98.63 |
| Thermoanaerobacter brockii | NC_014964.1 | 0.90 | 39 | - | - | - |
| Thermoanaerobacter sp | NC_014538.1 | 0.90 | 38 | 62.12 | - | 54.04 |
| Thermococcus sibiricus | NC_012883.1 | 0.80 | 50 | 99.00 | 95.95 | - |
| Variovorax paradoxus | NC_012792.1 | 0.80 | 82 | 98.44 | 99.48 | - |
| Weeksella virosa | CP002455.1 | 0.80 | 41 | 98.53 | - | 98.86 |
| Wolbachia sp | NC_012416.1 | 0.80 | 64 | 63.15 | - | 61.90 |
| Xanthobacter autotrophicus | NC_009720.1 | 0.70 | 17 | 90.11 | - | 90.35 |
| Yersinia pestis | NC_010159.1 | 0.40 | 20 | 59.47 | - | 91.93 |

Table 4: Information about the recovered species for the simHC+ dataset. "-" represents that the species was not recovered by the binning tool.

| Dataset | Available contigs to bin | No. of bins and contigs binned | MaxBin2 [6] | MetaBAT2 [2] | Vamb [3] | MetaCoAG |
|---|---|---|---|---|---|---|
| simHC+ [7] | 6,706 | Bins detected<br>Contigs binned | 95<br>5,723 | 32<br>54 | N/A<br>N/A | 90<br>6,033 |
| CAMI UG [4] | 49,927 | Bins detected<br>Contigs binned | 98<br>49,686 | 202<br>1,947 | 100<br>21,398 | 83<br>38,839 |
| CAMI Skin [4] | 139,217 | Bins detected<br>Contigs binned | 176<br>137,185 | 240<br>7,619 | 167<br>64,626 | 106<br>94,989 |
| CAMI Oral [4] | 96,079 | Bins detected<br>Contigs binned | 137<br>95,169 | 152<br>2,621 | 176<br>76,024 | 106<br>68,009 |
| CAMI GI [4] | 75,913 | Bins detected<br>Contigs binned | 127<br>75,509 | 389<br>6,659 | 156<br>41,948 | 113<br>57,251 |
| CAMI Airways [4] | 142,477 | Bins detected<br>Contigs binned | 155<br>140,415 | 205<br>10,924 | 173<br>69,353 | 96<br>101,732 |
| Sharon [5] | 7,067 | Bins detected<br>Contigs binned | 14<br>7,061 | 24<br>199 | 10<br>453 | 10<br>1,982 |
| COPD [1] | 67,753 | Bins detected<br>Contigs binned | 156<br>65,039 | 76<br>3,121 | 61<br>7,828 | 68<br>59,974 |

Table 5: The number of bins detected and the number of contigs binned by the binning tools for the co-assemblies of all the datasets.

| Dataset | Tool | Total number of bins detected | Number of high-quality bins | Number of medium-quality bins | Number of low-quality bins |
|---|---|---|---|---|---|
| simHC+ [7] | MaxBin2 | 95 | 59 | **11** | 25 |
| | MetaBAT2 | 32 | 4 | 4 | 24 |
| | MetaCoAG | 90 | **69** | 8 | **13** |
| CAMI UG [4] | MaxBin2 | 98 | 49 | **17** | 32 |
| | MetaBAT2 | 202 | 5 | 12 | 185 |
| | Vamb | 100 | 34 | 10 | 56 |
| | MetaCoAG | 83 | **50** | **17** | **16** |
| CAMI Skin [4] | MaxBin2 | 176 | 42 | 30 | 104 |
| | MetaBAT2 | 240 | 0 | 5 | 235 |
| | Vamb | 167 | 36 | 15 | 116 |
| | MetaCoAG | 106 | **49** | **33** | **24** |
| CAMI Oral [4] | MaxBin2 | 137 | 54 | **24** | 59 |
| | MetaBAT2 | 152 | 1 | 7 | 144 |
| | Vamb | 176 | 45 | 19 | 112 |
| | MetaCoAG | 106 | **58** | 19 | **29** |
| CAMI GI [4] | MaxBin2 | 127 | **59** | 22 | 46 |
| | MetaBAT2 | 389 | 4 | 9 | 376 |
| | Vamb | 156 | 44 | 14 | 98 |
| | MetaCoAG | 113 | 57 | **28** | **28** |
| CAMI Airways [4] | MaxBin2 | 155 | 32 | 26 | 97 |
| | MetaBAT2 | 205 | 1 | 2 | 202 |
| | Vamb | 173 | 20 | 14 | 139 |
| | MetaCoAG | 96 | **33** | **29** | **34** |
| Sharon [5] | MaxBin2 | 14 | 6 | 2 | 6 |
| | MetaBAT2 | 24 | 2 | 2 | 20 |
| | Vamb | 10 | 5 | 1 | 4 |
| | MetaCoAG | 10 | **7** | **3** | **0** |
| COPD [1] | MaxBin2 | 156 | 9 | 24 | 123 |
| | MetaBAT2 | 76 | 0 | 2 | 74 |
| | Vamb | 61 | 6 | 7 | 48 |
| | MetaCoAG | 68 | **17** | **25** | **26** |

Table 6: The number of high-quality, medium-quality and low-quality bins produced by each binning tool for the co-assemblies of all the datasets.

| Dataset | Tool | Running time (wall time) | Memory usage (kbytes) |
|---|---|---|---|
| simHC+ [7] | MaxBin2 | 5m 37s | 4,063, 345 |
| | MetaBAT2 | 12s | 619,345 |
| | MetaCoAG | 26m 34s | 621,584 |
| CAMI UG [4] | MaxBin2 | 26m 04s | 2,604,084 |
| | MetaBAT2 | 1m 51s | 985,860 |
| | Vamb | 2h 37m 32s | 701,764 |
| | MetaCoAG | 19m 24s | 1,944,632 |
| CAMI Skin [4] | MaxBin2 | 1h 58m 56s | 1,184,476 |
| | MetaBAT2 | 7m 21s | 2,185,080 |
| | Vamb | 8h 18m 42s | 976,644 |
| | MetaCoAG | 1h 01m 44s | 5,291,572 |
| CAMI Oral [4] | MaxBin2 | 1h 11m 49s | 1,130,420 |
| | MetaBAT2 | 3m 42s | 1,585,200 |
| | Vamb | 6h 31m 47s | 891,608 |
| | MetaCoAG | 43m 24s8 | 4,411,072 |
| CAMI GI [4] | MaxBin2 | 59m 53s | 1,874,764 |
| | MetaBAT2 | 2m 53s | 1,531,248 |
| | Vamb | 3h 26m 43s | 741,960 |
| | MetaCoAG | 32m 16s | 2,569,092 |
| CAMI Airways [4] | MaxBin2 | 1h 51m 35s | 1,338,220 |
| | MetaBAT2 | 6m 28s | 2,145,720 |
| | Vamb | 10h 04m 16s | 1,035,716 |
| | MetaCoAG | 1h 03m 13s | 6,476,888 |
| Sharon [5] | MaxBin2 | 1m 27s | 384,264 |
| | MetaBAT2 | 9s | 212,252 |
| | Vamb | 34m 20s | 607,708 |
| | MetaCoAG | 7s | 417,344 |
| COPD [1] | MaxBin2 | 1h 09m 17s | 352,888 |
| | MetaBAT2 | 1m 31s | 950,584 |
| | Vamb | 6h 06m 54s | 894,956 |
| | MetaCoAG | 13m 50s | 3,967,256 |

Table 7: Running time and memory usage of the different binning tools for all the datasets. All the binning tools were run on a Linux system with Ubuntu 18.04.1 LTS, 16 GB memory and Intel(R) Core(TM) i7-7700 CPU @ 3.60 GHz with 4 CPU cores. 8 threads were used for binning tools which support parallel execution.

[3] Jakob Nybo Nissen, Joachim Johansen, Rosa Lundbye Allesøe, Casper Kaae Sønderby, Jose Juan Almagro Armenteros, Christopher Heje Grønbech, Lars Juhl Jensen, Henrik Bjørn Nielsen, Thomas Nordahl Petersen, Ole Winther, and Simon Rasmussen. Improved metagenome binning and assembly using deep variational autoencoders. *Nature Biotechnology*, Jan 2021.

[4] Alexander Sczyrba, Peter Hofmann, Peter Belmann, David Koslicki, Stefan Janssen, et al. Critical Assessment of Metagenome Interpretation–a benchmark of metagenomics software. *Nature Methods*, 14:1063–1071, Oct 2017.

[5] Itai Sharon, Michael J. Morowitz, Brian C. Thomas, Elizabeth K. Costello, David A. Relman, and Jillian F. Banfield. Time series community genomics analysis reveals rapid shifts in bacterial species, strains, and phage during infant gut colonization. *Genome Research*, 23(1):111–120, 2013.

[6] Yu-Wei Wu, Blake A. Simmons, and Steven W. Singer. MaxBin 2.0: an automated binning algorithm to recover genomes from multiple metagenomic datasets. *Bioinformatics*, 32(4):605–607, Oct 2015.

[7] Yu-Wei Wu, Yung-Hsu Tang, Susannah G. Tringe, Blake A. Simmons, and Steven W. Singer. MaxBin: an automated binning method to recover individual genomes from metagenomes using an expectation-maximization algorithm. *Microbiome*, 2(1):26, Aug 2014.
